# Multiple Intermediates in the Detergent-Induced Fusion of Lipid Vesicles

**DOI:** 10.1101/2023.09.21.558601

**Authors:** Lara G. Dresser, Casper Kunstmann-Olsen, Donato Conteduca, Christopher M. Hofmair, Nathan Smith, Laura Clark, Steven Johnson, J. Carlos Penedo, Mark C. Leake, Steven D. Quinn

## Abstract

Detergent-induced vesicle interactions, critical for applications including virus inactivation, varies according to the detergent type and membrane composition, but the underlying mechanistic details remain underexplored. Here, we use a lipid mixing assay based on Förster resonance energy transfer (FRET), and single-vesicle characterization approaches to identify that sub-micron sized vesicles are induced to fuse by the non-ionic detergent Triton-X 100. We demonstrate that the process is a multi-step mechanism, characterized by discrete values of FRET efficiency between membrane-embedded fluorophores, and involves permeabilization, vesicle docking, hemi-fusion and full lipid mixing at sub-solubilizing detergent concentrations. We also dissect the kinetics of vesicle fusion to surface-tethered vesicles using a label-free quartz-crystal microbalance with dissipation monitoring approach, opening a platform for biotechnology applications. The presented strategies provide mechanistic insight into the dynamics of vesicle fusion and have implications for applications including drug delivery and sensor development where transport and manipulation of encapsulated cargo is essential.

## Introduction

Detergent-membrane interactions are critical for biotechnological applications including membrane-protein extraction^1, 2^, cell lysis^3^, targeted drug delivery^4^, and virus inactivation^5, 6^. In this context, the widely used non-ionic detergent Triton X-100 (TX-100) plays a key role by efficiently and effectively solubilizing membrane bilayers. Open questions regarding the interaction, which involves membrane fusion as a key component^7, 8, 9^, include elucidating the detergent’s precise molecular-level mode of action, its role in lipid-lipid interactions, and the nature and morphology of the fused species.

Thermodynamic approaches aimed at characterizing the TX-100-membrane interaction have typically used highly-controllable model-membrane systems such as supported bilayers or synthetic unilammelar vesicles (UVs) to unveil concentration requirements necessary for solubilization^10^, and to disentangle the influence of membrane phase, composition, charge, and detergent: lipid ratio on the interaction^11, 12, 13^. In this context, UVs are classified as small (SUVs), large (LUVs) or giant (GUVs) reflecting vesicle sizes of < 100 nm, 100 nm - 1 μm and > 1 μm respectively. GUVs in particular have played a key role in interrogating the TX-100 membrane interaction^14^ with measurements broadly confirming that TX-100 induces solubilization at concentrations above its critical micellar concentration (CMC). However, dynamic light scattering (DLS) and turbidity approaches have also revealed an intriguing growth in vesicle size under sub-solubilizing conditions^7, 8, 9, 15^, pointing towards morphological changes such as membrane swelling and fusion^8^ that could be harnessed for biotechnological applications.

To assess the swelling aspect, single-vesicle characterization approaches based on the fluorescence imaging of GUVs have revealed detergent-induced surface-area enhancements concurrent with the formation of dynamic pores^16, 17, 18^. Similar work also revealed that the membrane tension on both micron- and 100 nm-diameter vesicles decreases with TX-100 concentration, and the detergent introduces phase separation events in vesicles incorporating cholesterol^13, 18^. More recently, we demonstrated that variations in the efficiency of Förster resonance energy transfer (FRET) between fluorescently labelled lipids in LUVs quantitatively reports on their expansion upon TX-100 addition^19^, and combinations of molecular dynamics simulations and fluorescence approaches have revealed dynamic membrane bending events and the formation of local invaginations all prior to micellization^20^.

These results, and others^21^, are broadly supportive of a dynamic model for TX-100 induced membrane solubilization that involves detergent saturation on the membrane surface, the formation of mixed detergent-lipid micelles, structural changes within the intact membrane, and the release of mixed micelles to solution. A more quantitative extension of this model involves the formation of phase- and composition-sensitive pores, content leakage, variations in membrane curvature and local changes to the bilayer fluidity^22^. However, the molecular mechanisms underpinning another major class of event - namely TX-100 induced vesicle fusion - are not well characterized owing in part to a lack of experimental techniques capable of capturing the entire process.

In the context of vesicle fusion, optical microscopy experiments based on the imaging of unlabelled GUVs incubated with fluorescently tagged native membranes at sub-solubilizing TX-100 concentrations have indicated the presence of fusion by the appearance of fluorescence at the GUV site^23^. Using similar approaches, fusion between proteoliposomes and native membranes has been harnessed to enable reconstitution of transmembrane proteins into GUV bilayers after TX-100 addition^7, 8, 9, 24^. Phase contrast imaging and DLS measurements also indicate that the degree of fusion is dependent on the lipid composition, suggesting that the TX-100-membrane interaction may be a much more complex and multifaceted process. Confocal-based fluorescence experiments have also been employed to assess permeable GUV fusion, and although fusion was rarely observed, the results point towards a situation where fusion can occur due to casual contact between pores on adjacent membranes^17^. Unfortunately, conventional microscopy techniques are limited in their ability to resolve structural changes in objects smaller than 5-10 μm, can only quantify macroscopic changes in size and packing density and they lack the ability to follow molecular-level structural rearrangements on the nanoscale. There is therefore demand for alternative methods that can monitor the fusion of individual vesicles with sizes comparable to LUVs. In this regard, molecular dynamics simulations have alluded to a stock-pore mechanism of spontaneous bilayer fusion where initial vesicle docking precedes the formation of an hourglass-shaped connection between two adjoining membranes before hemi-fusion and full lipid mixing^17, 25, 26^. Whether such a mechanism occurs in detergent-rich media, and whether these states can be controllably harnessed remains an open question.

Inspired by these insights and in search of the mechanistic origins of TX-100 induced vesicle fusion, we adopted a range of ensemble- and single-vesicle characterization approaches to interrogate the structural integrity of highly curved LUVs composed of POPC (1-palmitoyl-2-oleoyl-glycero-3-phosphocholine) in response to TX-100. The LUVs used here differ from low-curvature GUV species by displaying a maximal principal curvature^27^ of ∼10^7^m^-1^, which is an order of magnitude greater than those typically observed in GUVs. By employing particle sizing approaches including DLS and fluorescence correlation spectroscopy (FCS), we first confirm an enhancement in vesicle size under sub-solubilizing conditions. We next showcase a steady-state and time-resolved assay based on FRET^28, 29^, to quantitatively assess the degree of TX-100 induced lipid mixing between vesicles. Based on the magnitude of the observed FRET efficiency between fused vesicles, coupled with morphological changes observed via electron microscopy approaches, we provide evidence that the detergent-induced fusion process is a time-dependent and multi-step process comprising vesicle docking, hemifusion and full lipid mixing prior to solubilization. While we have previously used FRET-based methods to quantitatively report on vesicle swelling^19, 30, 31^, here we take advantage of enhancements in the FRET efficiency to quantify the magnitude and state of fusion. To further characterize the interaction, we also deploy a calcium-sensing assay to confirm TX-100 induces vesicle permeabilization during the process. Finally, to demonstrate the potential of TX-100 to facilitate vesicle fusion of freely-diffusing vesicles to surface-immobilized vesicles, and to extract kinetic details of the process, we deployed an acoustic-sensing approach based on quartz-crystal microbalance with dissipation monitoring (QCM-D). In this regard, the QCM-D approach offers highly sensitive real-time monitoring of surface mass and viscoelasticity, without the need for fluorescent labels. This not only opens exciting possibilities for the controllable delivery of encapsulated cargo to a substrate but may also support a range of bionanotechnological applications including diagnostic sensing, drug delivery and the formation of biomaterials.

Our discovery of a multi-step and controllable fusion process in both freely diffusing and surface-tethered vesicles at sub-solubilizing detergent concentrations provides clues to the underlying TX-100-membrane interaction and may have important implications for the transport of membrane proteins such as proton pumps and transport receptors^23^. We also expect the presented techniques to have general applicability for assessing interactions between highly curved lipid vesicles and a wide range of fusogens, such as neurotoxic proteins with important biomedical significance.

## Results

### Triton X-100 Alters the Morphology of Vesicles at Sub-Solubilizing Concentrations

LUVs composed of POPC lipids were prepared by the extrusion method and confirmed to be ∼200 nm in diameter by DLS (**Supplementary Figure 1**). We first employed FCS and DLS to assess morphological changes as TX-100 was progressively added to a solution containing freely diffusing vesicles. Both techniques are sensitive to the diffusion coefficient of vesicles in solution, and thus quantitatively report on their hydrodynamic radius via the Stokes-Einstein relationship^32^. The FCS correlation curves associated with freely diffusing LUVs labelled with 0.1 % of the fluorescent membrane dye DiI progressively shifted towards longer lag times at detergent concentrations below the reported CMC (∼0.2-0.3 mM)^3, 33^ (**Figure 1a**) - a finding that warranted further attention given non-ionic surfactants typically only achieve membrane solubilization once above the CMC. Here, a reduction of 40 % in vesicle diffusion coefficient, from an initial starting value of 1.28 ± 0.01 μm^2^s^-1^ (±SD) in the absence of TX-100, was observed, corresponding to an increase in hydrodynamic diameter (d_H_) from 168 ± 1 nm to 290 ± 5 nm. In all cases, the FCS curves were best fitted to a single diffusing component model (see **Methods**) with low residuals, and notably lacked a significant population of micelles exhibiting higher diffusion coefficients. In line with previous work^19^, we attributed these observations to vesicle swelling, docking, fusion, aggregation, or combinations of these effects in solution. At TX-100 concentrations beyond the CMC, the particle radius reduced to ∼3.5 nm which we assigned to solubilization and the formation of mixed detergent-lipid micelles. DLS also confirmed a positive shift in size distribution, though we note in these experiments, the lipid concentration was increased by two orders of magnitude relative to the FCS work, which we hypothesised could facilitate fusion. Here, correlation curves obtained from the intensity of scattered light also progressively shifted towards longer lag times (**Figure 1b**). Without detergent, the vesicles displayed a lognormal size distribution, as expected for freely diffusing species, peaking at 182 nm with a low polydispersity index of 0.16 ± 0.01 (**Figure 1c, Supplementary Figure 2**). At detergent concentrations approaching the CMC, the correlation curves then revealed a 1.7-fold increase in vesicle size (**Figure 1c**) and ∼4-fold increase in polydispersity index (**Figure S2**), suggestive of mixed species in the ensemble. At 0.5 mM TX-100, most particles displayed d_H_ in the range 6 - 10 nm, consistent with mixed detergent-lipid micelles. However, we also observed objects with sizes ranging from 250 - 480 nm, in line with expanded, fused and/or aggregated species, and to a lesser extent, particles with intermediate diameters, which could be indicative of bicelles and cylindrical micelles^34^ (**Figure S2**). When intensity distributions were converted into volume distributions, similar trends were observed (**Figure S2**). Caution should be taken in interpretating such distributions due to underlying assumptions related to vesicle morphology and noise amplification, but below the CMC, the distributions broadened with TX-100 addition, and a mixture of species was exposed at 0.5 mM, in line with our intensity-based analysis. While a direct comparison between FCS and DLS datasets is not entirely straightforward owing to variations in the initial lipid concentration, both analyses are broadly supportive of a progressive increase in particle size below the CMC.

**Figure 1.**
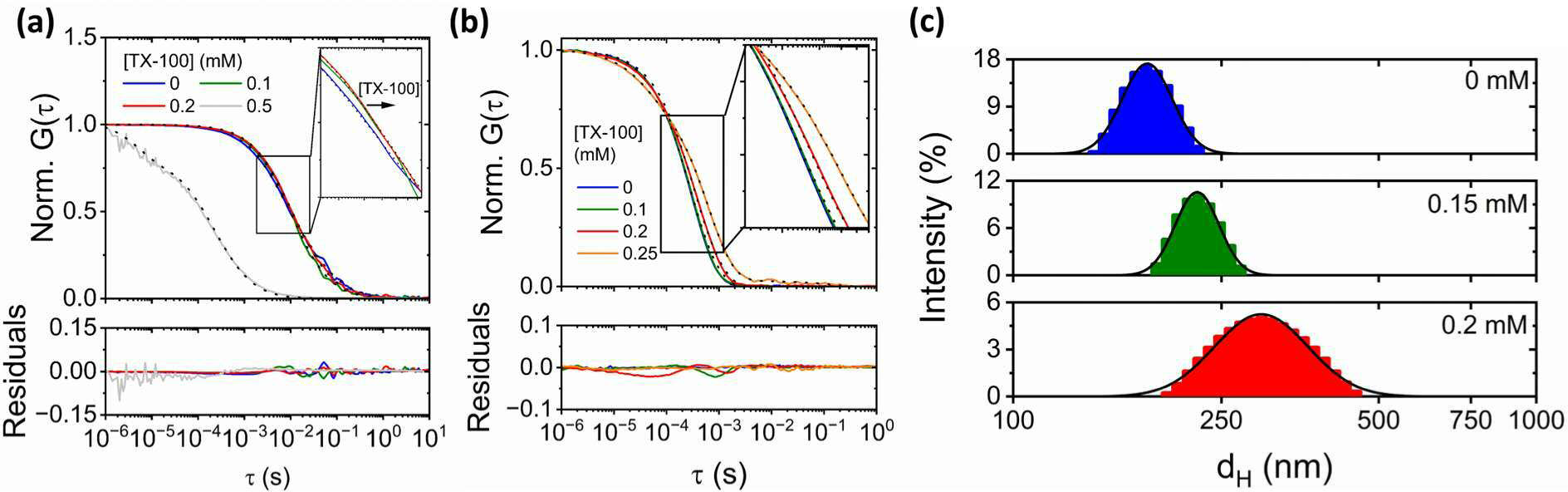
Characterization of vesicle-detergent interactions at sub-solubilizing TX-100 concentrations. (a) Top: Normalized variation in FCS correlation curves (solid lines) and fits (dotted lines) associated with interactions between LUVs and 0 (blue), 0.1 (green), 0.2 (red) and 0.5 (grey) mM TX-100 at 21°C. Bottom: corresponding residuals of the fits. (b) Top: Normalized variation in DLS correlation curves (solid lines) and fits (dotted lines) associated with interactions between LUVs and 0 (blue), 0.1 (green), 0.2 (red) and 0.25 (orange) mM TX-100 at 21°C. Bottom: residuals of the fits. (c) Corresponding hydrodynamic diameter distributions obtained from LUVs in the absence (blue) and presence of 0.15 (green) and 0.2 (red) mM TX-100. The solid black lines represent lognormal fits with peak d_H_ values of 182.6 ± 0.1 nm (Chi-squared χ_r_^2^ = 0.02) (top), 226.8 ± 0.2 nm (χ_r_^2^ = 0.02) (middle) and 311.2 ± 0.7 nm (χ_r_^2^ = 0.01) (bottom), respectively.

### Triton X-100 Induces Vesicle Fusion

To directly test for fusion, we next employed a lipid mixing assay to monitor interactions between LUVs containing 2 % DiI (FRET donor) and those containing 2 % DiD (FRET acceptor) (**Figure 2a**). Direct excitation of DiI in the absence of fusion thus yields low sensitized emission from the acceptor, however, upon vesicle fusion, lipid mixing brings the FRET pairs into close proximity, triggering enhanced sensitized acceptor emission as a consequence of non-radiative energy transfer^28, 29, 35^. A key benefit of the approach is that the total number of fluorophores remains constant throughout the fusion process, allowing for intermediate states, corresponding to different levels of lipid mixing, to be observed by quantifiable variations in the FRET efficiency (Equation 2). Ensemble FRET measurements were performed as an initial step to quantify interactions between vesicles containing DiI and DiD in response to TX-100. Here, we measured the efficiency of energy transfer using the *RatioA* parameter (Equation 3) which compares the amount of sensitized acceptor emission with that obtained by direct excitation of the acceptor. In the absence of TX-100 the intensity of sensitized acceptor emission obtained from a 1: 3 mixture of DiI: DiD vesicles was minimal, then progressively increased upon TX-100 addition towards the CMC and was anti-correlated with quenching of the donor (**Figure 2b**, **Supplementary Figure 3**). The anti-correlated nature of the DiI and DiD signals represents the classic hallmark of FRET^36^, providing assurance that the observed changes are due to energy transfer. Beyond the CMC, the apparent FRET efficiency then decreased towards its initial value, reflected by an enhancement of the donor signal and corresponding reduction in sensitized acceptor emission (**Figure 2b, Supplementary Figure 3**). The increase in *RatioA* observed below the CMC indicates a progressive decrease in the mean donor-acceptor distance, supporting a model where progressive addition of TX-100 shifts the vesicle population from intact to fused species prior to solubilization and the formation of mixed micelles (**Figure 2c**). Indeed, this is reflected in the *RatioA* response, where a progressive increase towards a peak of 0.49 ± 0.03 at the CMC was observed, suggestive of morphological changes in the ensemble that results in close proximity of the dyes, followed by a reduction towards its initial value, indicative of solubilization (**Figure 2d**). To provide confidence that the observed increase in *RatioA* was due to lipid mixing, and not lipid exchange between separated vesicles, we also assessed the response of the labelled vesicles in the presence of various cyclodextrins. As previously highlighted, methyl-β-cyclodextrin (MβCD) and 2-hydropropyl-β-cyclodextrin (HPβCD) induce lipid exchange at sub-solubilizing concentrations^37^, and both were added here to a solution of vesicles identical to those shown in **Figure 2b**. We observed that the sensitized acceptor emission (and corresponding *RatioA* and *E_FRET_* signals) increased slightly above the baseline (*E_FRET_* ∼ 0.09) upon both MβCD and HPβCD addition (**Supplementary Figure 4**), and that the FRET signal was enhanced when DiI and DiD were replaced with Lipo-Cy3-CO and Lipo-Cy5-N_3_ (**Supplementary Figure 4**). As previously reported, the Lipo derivatives are structurally similar to DiI and DiD (**Supplementary Figure 4**), but their FRET is enhanced upon cycloaddition^38^. We interpreted the combined data as evidence that both cyclodextrins can, to some extent, exchange DiI and DiD, however, this may not be as efficient as the exchange of unlabelled lipids. To explicitly test for DiI and DiD exchange, we also performed an experiment whereby 200 nm sized POPC vesicles labelled with 2 % DiI were separated from vesicles containing 2 % DiD by a dialysis membrane (MWCO = 3 kDa) (**Supplementary Figure 5**). In this case, we prevented DiI and DiD labelled vesicles from directly interacting, but allowed low molecular weight species, including free lipids, to diffuse across the membrane (**Supplementary Figure 5**). However, after 5 hours of dialysis in the absence and presence of TX-100, we observed no evidence of DiD absorption or sensitized emission after observing the solution containing DiI. We did, however, observe increased levels of scattering upon TX-100 addition, which we interpreted as a signature of fusion. Taking the combined data together, the implication is that the data shown in **Figure 2b** is more likely due to DiI and DiD mixing as opposed to DiI and DiD exchange.

**Figure 2.**
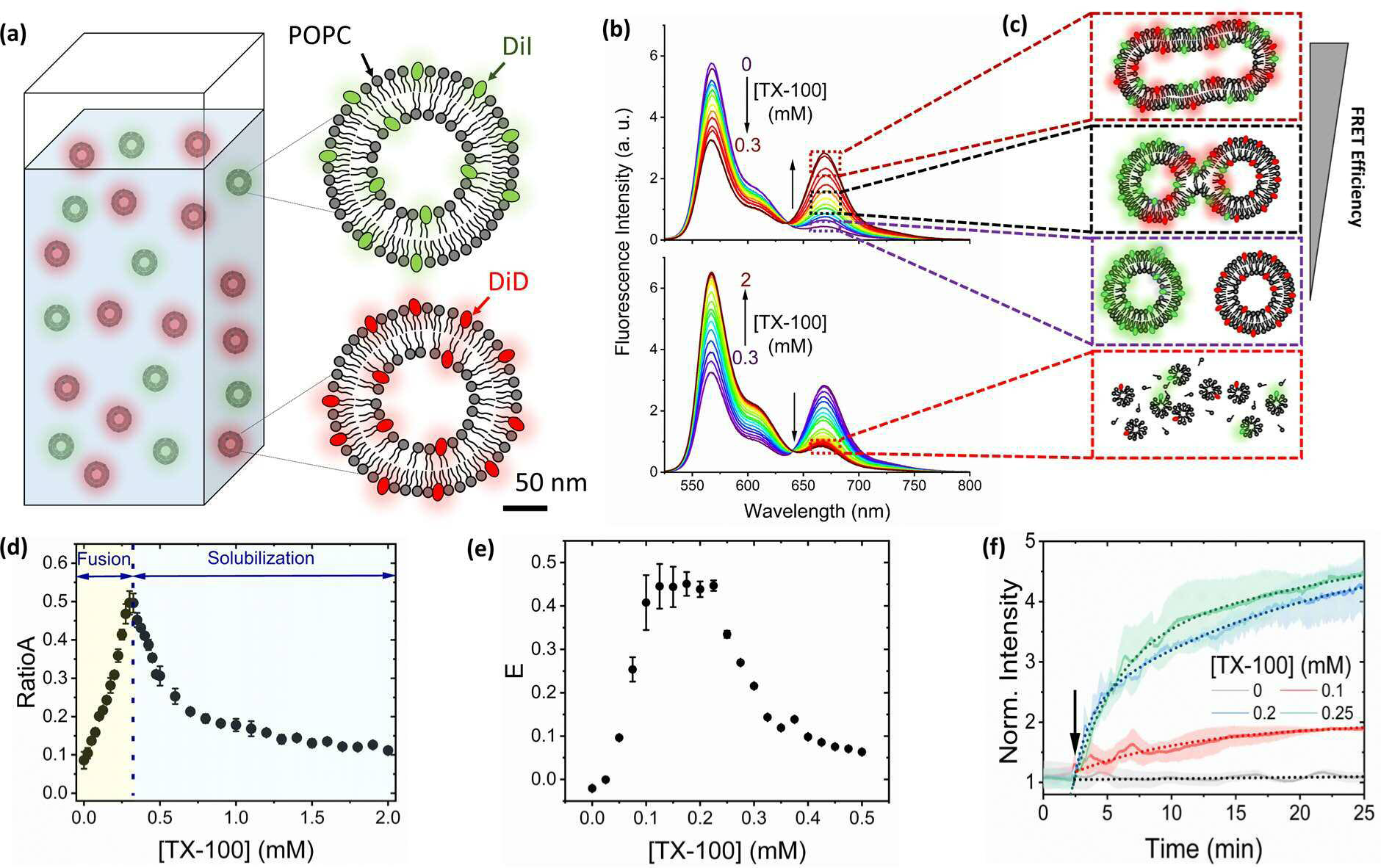
TX-100 induces vesicle fusion. (a) Schematic illustration of the lipid mixing assay. POPC LUVs containing 2 % DiI (green) and 2 % DiD (red) are incubated at a 1: 3 ratio. (b) Representative variation in fluorescence emission spectra (λ_ex_ = 520 nm) obtained from LUVs in the presence of 0 - 0.3 mM (top panel) and 0.3 mM - 2 mM TX-100 (bottom panel) at 21°C. (c) Mixing LUVs with TX-100 results in FRET enhancements assigned to vesicle docking and fusion prior to FRET reductions attributed to micellization. (d) Corresponding variation in *RatioA*. (e) Variation in FRET efficiency, *E*, as estimated from changes in τ*_D_* and τ*_DA_* across the titration. Data points represent the mean and standard error of the mean from three separated experimental runs. (f) Time-dependent variation in DiD emission upon injection of TX-100 at 21°C (λ_ex_ = 520 nm). The grey dashed line corresponds to a linear fit, coloured dashed lines correspond to bi-exponential fits and the shaded areas correspond to the SE. The arrow indicates the point of TX-100 injection.

When additional control experiments involving the addition of TX-100 to a 1:3 mixture of LUVs lacking fluorophores and those containing 2 % DiD were performed, similar results were obtained (**Supplementary Figure 6**), indicating further that the observed increase in *RatioA* shown in **Figure 2b** is due to energy transfer and sensitized acceptor emission, as opposed to photophysical phenomenon such as fluorescence de-quenching. We also note that the amounts of DiI and DiD per vesicle and the DiI: DiD ratio were optimized (1: 3, 2 % of each dye) to maximize the magnitude of the FRET response across the interaction (**Supplementary Figure 7**).

To confirm the presence of an energy transfer mechanism between DiI and DiD during the interaction, we also evaluated the amplitude-weighted fluorescence lifetime of DiI in the absence, τ*_D_*, and presence, τ*_DA_*, of DiD. In all cases, the lifetime decays displayed bi-exponential behaviour after deconvolution with the instrument response function (**Supplementary Figure 8**) (Equation 4). As TX-100 was progressively titrated at concentrations below the CMC, τ*_D_* progressively increased from an initially self-quenched value of 0.53 ± 0.03 ns to 1.36 ± 0.02 ns, representing the unquenched DiI lifetime (**Supplementary Figure 9**).We also found that τ*_DA_* remained largely unchanged from its initial starting value of ∼0.54 ns, then sharply increased towards ∼1.2 ns as the concentration of TX-100 increased beyond the CMC (**Supplementary Figure 9**). We note that τ*_D_* and τ*_DA_* displayed similar initial starting values which we interpreted as evidence of minimal spontaneous fusion prior to injection of TX-100. The corresponding variation in the efficiency of energy transfer, *E* (Equation 5), which takes into account variations in τ*_DA_* relative to τ*_D_*, then displayed several common features to the *RatioA* measurements, including (i) a progressive increase as the TX-100 concentration approached the CMC, (ii) a turning point at the CMC and (iii) a progressive reduction towards its initial value during the solubilization step (**Figure 2e**). The observed increase in *RatioA* and *E*, which effectively remove contributions from DiI de-quenching, provide confidence of a TX-100-induced solubilization mechanism comprising vesicle fusion (high FRET states) below the CMC, followed by solubilization and mixed micelle formation above the CMC (low FRET states) (**Figure 1c**).

We also note the time-dependent increase in sensitized DiD emission following TX-100 injection followed bi-exponential kinetics, pointing further towards a fusion process involving multiple stages. In the absence of TX-100, DiD emission remained largely invariant across the observation time window, also indicating a lack of spontaneous fusion, however, upon TX-100 addition, bi-exponential kinetics were monitored where the amplitude weighted time constant was 273 ± 6 s in the presence of 0.1 mM TX-100, and 396 ± 11 s in the presence of 0.25 mM (**Figure 2f, Supplementary Table 1**). While the precise nature of the bi-exponential kinetics required further exploration, we hypothesised that the various components are more likely to represent fusion steps (for example hemi-fusion and full lipid mixing) versus combinations of lipid exchange and fusion.

To further characterize the interaction and assess the degree of membrane permeabilization and solution exchange during the TX-100 LUV interaction, we also implemented an approach based on the measurement of Ca^2+^ entry into vesicles encapsulating the fluorescent calcium indicator Cal-520. This approach is similar in nature to other permeabilization assays^30, 39^ whereby Ca^2+^ flux into intact, surface-tethered vesicles triggers a concentration dependent intensity enhancement of Cal-520 at ∼520 nm, as illustrated in **Supplementary Figure 10**. Prior to experimentation, nonencapsulated molecules were removed by size exclusion chromatography as detailed in the **Methods**. Immobilized vesicles were then imaged in buffer solution containing 10 mM Ca^2+^ with and without addition of TX-100. We imaged multiple fields of view per condition, enabling quantification of intensity distributions from several thousand vesicles before adding buffer solution containing TX-100 and Ca^2+^. In the absence of TX-100 and Ca^2+^, we identified ∼100 vesicles per 25 x 50 μm field of view, and the intensity distribution fitted to a log-normal model with a peak position of 135 counts / 100 ms. When 10 mM Ca^2+^ was flushed across the same vesicles, the number of fluorescent foci remained invariant, and the intensity distribution was comparable, peaking at 147 counts/ 100 ms. A similar dataset was observed when the vesicles were incubated with 0.025 mM TX-100 and 10 mM Ca^2+^, however in the presence of 0.05 - 0.1 mM TX-100, we observed brighter foci and clearly discernible shifts in the distributions from low- to-high intensities. In such cases, a 2-fold or greater increase in the probe’s emission intensity was observed, in line with those previously identified^30, 40^. Beyond 0.1 mM, the number of spots decreased ∼10-fold and each had an intensity below the baseline level, indicating Cal-520 leakage from the LUVs. Control experiments performed independently on free Cal-520 in solution indicated that the dye’s fluorescence is insensitive to TX-100, and that it responds to Ca^2+^ in detergent-rich media (**Supplementary Figure 11**), providing confidence that the observed intensity enhancements are due to membrane permeabilization and Ca^2+^ transport across the membrane. Importantly, the number of surface-immobilized vesicles on the surface remained largely unchanged as TX-100 up to 0.1 mM was introduced. When compared with the FRET measurements shown in **Figure 2d and 2e**, which indicate substantial levels of lipid mixing at comparable TX-100 concentrations, the implication is that combinations of membrane permeabilization and content exchange occur concurrently with fusion events in the ensemble. However, whether content can be controllably recovered after release from the vesicles during the permeabilization and/or solubilization stages remains an open question.

### The Fusion Mechanism involves Docking, Hemifusion and Full Lipid Mixing

To further explore the observed structural changes, and to dissect the mechanism of fusion, we next performed single-vesicle imaging via a custom-built wide-field objective-type total internal reflection fluorescence microscope that enables the simultaneous imaging of DiI and DiD emission^41^. Here, POPC vesicles incorporating 2 % DiI were induced to fuse with LUVs incorporating 2 % DiD and 1 % biotin-PE. Fused species were then tethered to a glass coverslip functionalized with biotinylated BSA and NeutrAvidin (**Figure 3a**). Vesicles were added to a surface-functionalized coverslip after incubation with TX-100, and DiI and DiD fluorescence trajectories were collected with 50 ms time integration. Fused vesicles were then identified by the appearance of co-localized diffraction limited spots across both detection channels (**Figure 3b**) and lipid mixing between vesicles was observed via an appearance or enhancement in the mean FRET efficiency per vesicle. In the absence and presence of 0 - 0.2 mM TX-100, the traces obtained from > 500 individual FRET-active species were photostable over the duration of the measurement (**Figure 3c, Supplementary Figure 12)** and the population distributions of FRET efficiencies showed a major peak at ∼0.1 (**Figure 3d**), suggestive of close contact or docking between DiI and DiD-labelled vesicles, but without substantial lipid mixing. As suggested elsewhere, a combination of factors, including Van der Waals, electrostatic and hydrophobic interactions may be responsible for some spontaneous vesicle-vesicle interactions, even in the absence of TX-100^42, 43^. As the TX-100 concentration then increased towards 0.5 mM, the traces remained photostable (**Supplementary Figure 12**) and the low FRET population decreased in favour of a broad distribution of high FRET states that reached a value of 0.7 at 0.5 mM, suggestive of substantial lipid mixing events (**Figure 3d**). We note the observed shift in the distributions from low to high FRET efficiencies broadly reflect the changes observed in the ensemble and the observation of FRET states between 0.1 and 0.7 reflect the morphological heterogeneity of intermediates that are structurally different from the fully fused state in which both leaflets have fully mixed. We also note that as the TX-100 concentration progressively increased across the titration, the total intensity per vesicle, defined as the sum of DiI and DiD emission intensities, increased > 2-fold from 1325 ± 75 counts/ 50 ms at 0.1 mM TX-100 to 3070 ± 182 counts / 50 ms. Since the particle size scales with the number of fluorophores^44^, the data shown in **Figure 3d** arises from the presence of large immobilized fusion products, as opposed to smaller mixed micelles. We therefore assigned *E_FRET_* = 0.7 as an indicator of full fusion, in which both leaflets of the vesicle fully mixed, whereas intermediate states were assigned to hemi-fusion, where only partial lipid mixing was achieved. In all cases, tri-Gaussian functions were fitted to the FRET efficiency histograms, and used to identify thresholds of *E_FRET_* < 0.25, 0.25 ≥ *E_FRET_* ≤ 0.55 and *E_FRET_* > 0.55 for classification of docked, hemi-fused and fully fused species respectively. As shown in **Figure 3e**, the percentage of species exhibiting full fusion increased 5-fold across the titration, whereas the number of docked species progressively diminished. While the current experiments do not inform on whether TX-100 promotes vesicle docking, its insertion into intact vesicles, coupled with its hydrophobicity and ability to form mixed detergent-lipid regions, could trigger vesicle-vesicle interactions via a route that involves vesicle destabilization as they dynamically interact^15^. Nevertheless, the imaging of single fused species via FRET provides evidence for the presence of intermediate states along the fusion process, with each state characterized by discrete values of FRET efficiency and thus DiI-DiD separation distance.

**Figure 3.**
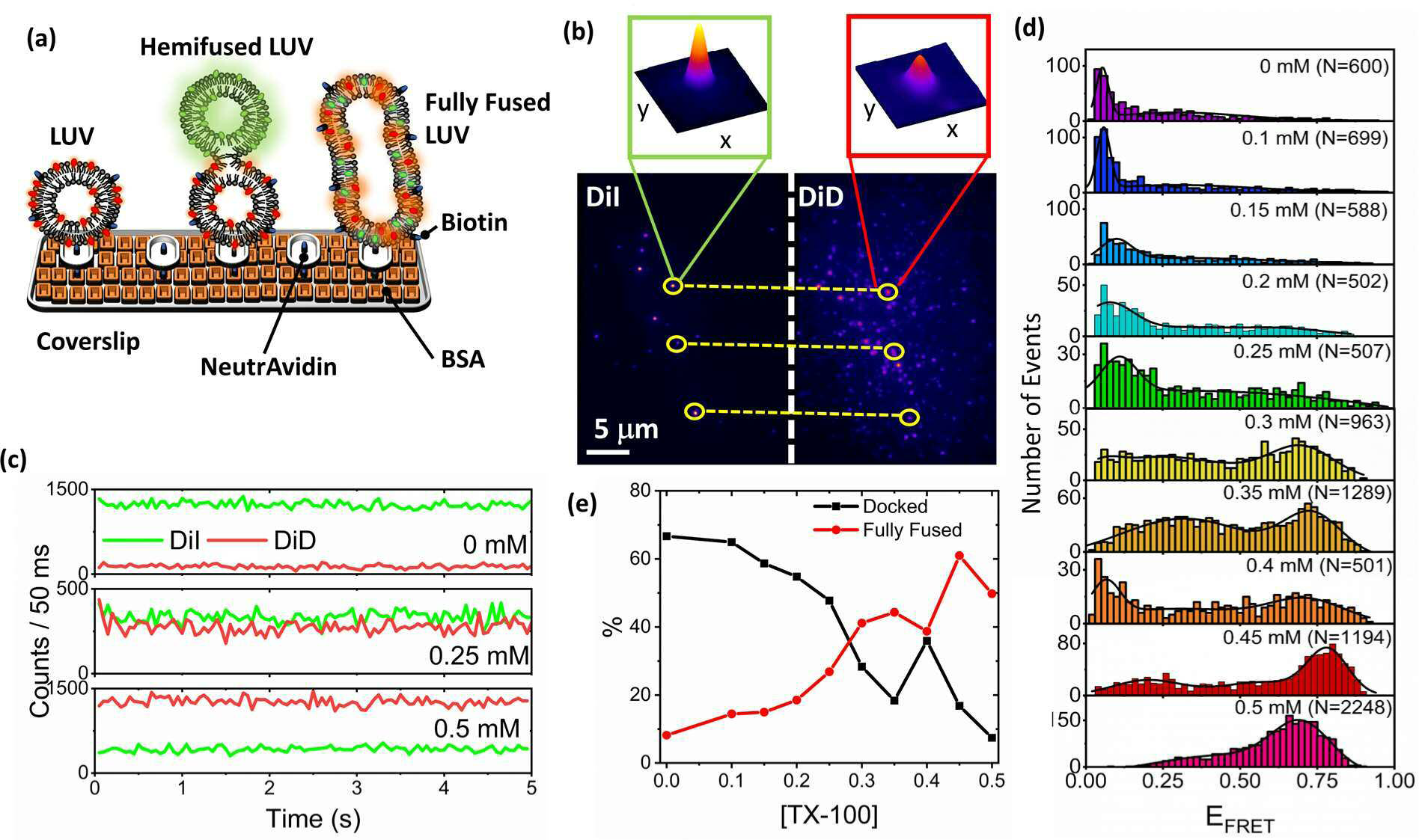
TX-100 induces vesicle docking, hemifusion and full lipid mixing. (a) Schematic illustration of the immobilization scheme. LUVs incorporating 2 % DiD and 1 % Biotin-PE were fused with LUVs incorporating 2 % DiI at 21°C and attached to a coverslip via BSA, biotin and NeutrAvidin. (b) Representative TIRF image obtained from surface-tethered LUVs in the presence of TX-100, showing co-localized DiI and DiD regions (yellow circles) (λ_ex_ = 532 nm). Insets: 3D intensity profiles of DiI and DiD emission obtained from a single fusion product. (c) Representative DiI (green) and DiD (red) intensity traces obtained from individually fused LUVs in the absence and presence of TX-100. (d) Histograms of the mean apparent FRET efficiency obtained from N > 500 surface immobilized vesicles before and after incubation with TX-100 rich solutions. Solid black lines represent tri-Gaussian fits. (e) Variation in the number of docked (*E_FRET_* < 0.25) versus fully fused (*E_FRET_* > 0.55) species observed as a function of TX-100.

To provide further evidence for the presence of multiple intermediates, we also acquired scanning electron microscopy (SEM) micrographs of vesicles in the absence and presence of fusion-promoting detergent concentrations. Without TX-100, the vesicles (N = 242) were predominantly spherical, exhibiting a mean circularity, ϕ, of 0.84 ± 0.15, and a mean particle size, d, of 246 ± 61 nm, with little evidence of fusion (**Figure 4a**, **Figure 4f, Supplementary Figure 13**). Upon incubation with TX-100, the mean particle size then increased 2-fold (d = 528 ± 56), the majority of species were elliptical (ϕ = 0.62 ± 0.22) (**Figure 4f)** and four major species were identified. First, we observed adjoined vesicles without evidence of clefting, which we interpreted as early fusion products that involve vesicle docking (**Figure 4b, Supplementary Figure 14**). Further examination revealed vesicles within close proximity but with a clear cleft between adjacent surfaces (**Figure 4c, Supplementary Figure 15**). Based on the similarity of these structures to those observed in previous studies of protein-induced vesicle fusion^45, 46^, we assigned these events to intermediate hemi-fusion products involving two or more vesicles. Finally, we observed the appearance of much larger micron-sized ellipses (**Figure 4d, Supplementary Figure 16**) and larger conglomerates (**Figure 4e, Supplementary Figure 17**) which we interpreted as evidence of multiple fusion events. While we previously identified that TX-100 leads to a 20 % increase in vesicle size through swelling^19^, the ellipses observed here were more than double the size of a single vesicle, and are therefore more likely to represent fully fused species as opposed to single expanded vesicles. Closer inspection of the surface topography of such species also indicated variations in texture, protrusions, and indentations which we speculate could be indicative of pore formation and/or dissociation of membrane regions (**Figure 4c**).

**Figure 4.**
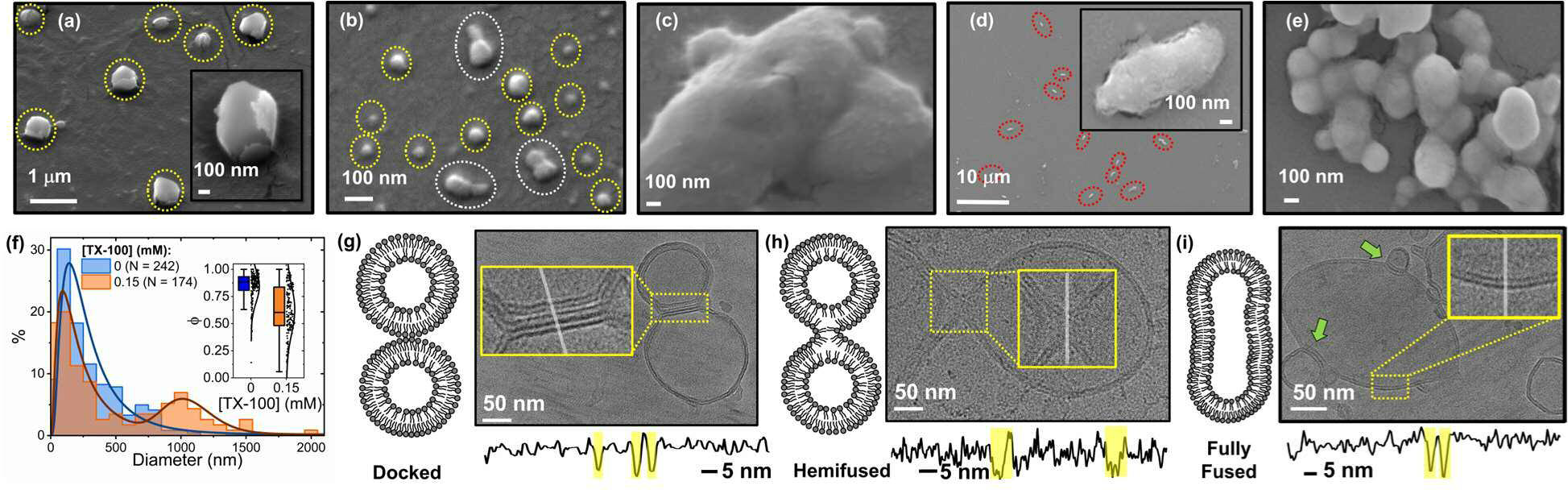
Confirmation of fused vesicles by SEM and Cryo-TEM. (a) SEM micrographs of intact vesicles in the absence and (b-e) presence of 0.15 mM TX-100 showing intact immobilized vesicles (yellow dashed regions), docked vesicles (white dashed regions), and large elliptical species (red dashed regions) assigned to full fusion. Vesicles were prepared and incubated with TX-100 at 21°C. (f) Size distribution histograms obtained from vesicles in the absence (blue) and presence (orange) of 0.15 mM TX-100. Inset: comparative box plots summarizing the corresponding variation in particle circularity. The boxes represent the interquartile ranges, the upper and lower whiskers represent the 5^th^ and 95^th^ percentiles, and the solid lines inside the boxes represent the medians. (g) Schematic (left panel) and representative cryo-TEM image (right panel) of docked, (h) hemi-fused and (i) fully fused species. Insets: regions of interest at higher magnification showing bilayers defined by the luminal grey level distribution (bottom panels). The green arrows indicate further docking or budding events.

Having demonstrated TX-100 induced morphological changes by SEM, we next used Cryo-TEM to visualize interactions between adjacent membranes. Here, the majority of isolated vesicles were intact and unilamellar, and in line with our prior SEM measurements, they exhibited a mean particle size of 249 ± 78 nm and were predominantly spherical (ϕ = 0.95 ± 0.04) (**Supplementary Figure 18**). In agreement with previous observations^47^, we also identified lower populations (< 40 %) of vesicles containing multiple bilayers (**Supplementary Figure 19**). Based on the prior FRET and SEM analysis, and in line with models predicted for extracellular vesicle fusion^48^, we hypothesized that TX-100 induced fusion intermediates consist of (i) docked vesicles showing close contact between the lipid bilayers, but without lipid mixing, (ii) progressive fusion of the outer and inner leaflets leading to a fusion pore and (iii) development of the fused pore into a fully fused state where both leaflets had fully mixed. Indeed, we identified examples of where the membranes of opposing vesicles were in parallel and within close proximity (< 5 nm), but with a clear barrier separating the vesicle interiors (**Figure 4g**). We also identified structures where the bilayer leaflets had merged at a ∼25 nm diameter contact pore, and where the two bilayers transitioned continuously from one vesicle to the other (**Figure 4h**). The fully fused species, after expansion of the fusion pore and upon complete content mixing, was then captured as a larger fusion product containing a single bilayer, with no evidence of a barrier preventing content mixing (**Figure 4i**). In some cases, we also observed examples of where further vesicles had either docked to or budded from the fully fused form (**Figure 4i**). Those species classified as docked, hemifused, or fully fused comprised 31 % of the population in 0.15 mM TX-100 and had a mean calliper diameter of 568 ± 86 nm, which we note represents a 2.3-fold increase relative to the intact vesicles (**Supplementary Figure 18**). The intermediates also displayed a shift towards lower circularity values, consistent with a global morphological change from spherical intact vesicles to larger, elliptical fusion products (**Supplementary Figure 18**). While the current Cryo TEM analysis cannot differentiate between further docking or budding events, we note that the smaller vesicles are connected to the main vesicle by a narrow aqueous channel pointing towards docked species. TX-100 has, however, been shown to induce vesicle budding in GUVs of similar composition^49, 50, 51^, and the release of smaller species cannot be ruled out. Nevertheless, these results further demonstrate the heterogeneous nature of the TX-100 induced fusion products.

### TX-100 Facilitates Fusion between Freely-Diffusing and Surface-Immobilized Vesicles

Controllable vesicle fusion to surface-tethered vesicles has vast potential for a range of applications including biosensing and drug delivery^52^. We therefore explored whether TX-100 could controllably fuse vesicles in solution with vesicles pre-immobilized on a solid surface, allowing for real-time content delivery and solution exchange. In contrast to our TIRF measurements, here we assessed the kinetics of accumulated mass transfer and variations in surface viscoelasticity as vesicles from solution interacted with those already pre-immobilized to a substrate via QCM-D. POPC vesicles incorporating 1 % biotin-PE were first tethered to a bovine serum albumin coated SiO_2_ substrate incorporating 2 % biotin via NeutrAvidin as described in **Figure 3a**. The frequency and dissipation signals, reflecting surface mass and viscoelasticity, respectively, were then recorded under equilibrium and exposure to a constant flow rate. **Supplementary Figure 20** shows representative QCM-D kinetics recorded upon injection of BSA-Biotin, Neutravidin and biotinylated vesicles. After the sensors were saturated with vesicles, the surface was washed with 50 mM Tris (pH 8) buffer to remove unattached species. Importantly, the frequency and dissipation responses from the sensor showed little recover after the wash step indicating successful surface tethering. Next, vesicles lacking biotin-PE at a final lipid concentration of 0.16 mg/mL were flushed across the sensor surface in the absence and presence of TX-100. Negligible vesicle deposition was noted in the absence of TX-100, indicated by the lack of variation in both the frequency and dissipation responses (**Supplementary Figure 20**). When a solution containing LUVs and 0.1 mM TX-100 was injected across the layer, an initial 20 Hz reduction in frequency, concurrent with a 1.2-fold increase in dissipation was recorded, indicative of (i) a mass gain over the period of the injection lasting tens of minutes and (ii) structural rearrangements on the surface (**Figure 5a**). We assigned the bi-exponential nature of the frequency decay to combinations of detergent interactions with the immobilized layer and vesicle-vesicle interactions on the surface; however, the QCM-D traces are unable to clearly distinguish between docked, hemi-fused and fully fused species. Experiments performed independently to assess the interaction between freely diffusing TX-100 with surface-tethered vesicles indicated only a 4-5 Hz shift upon injection (**Supplementary Figure 21**)^19^, consistent with a small mass gain on the sensor surface which we attribute to TX-100 incorporation into the vesicle bilayers until saturation is reached^19^. While the QCM-D data does not report on TX-100 flip-flop across the bilayer, we expect this to occur rapidly, and simultaneously with its incorporation^15^. Irrespective, the small shift observed here, indicates that the observed frequency change shown in **Figure 5a** is mostly due to TX-100 induced vesicle-vesicle interactions on the surface, as opposed to TX-100 incorporation into the surface-immobilized vesicles.

**Figure 5.**
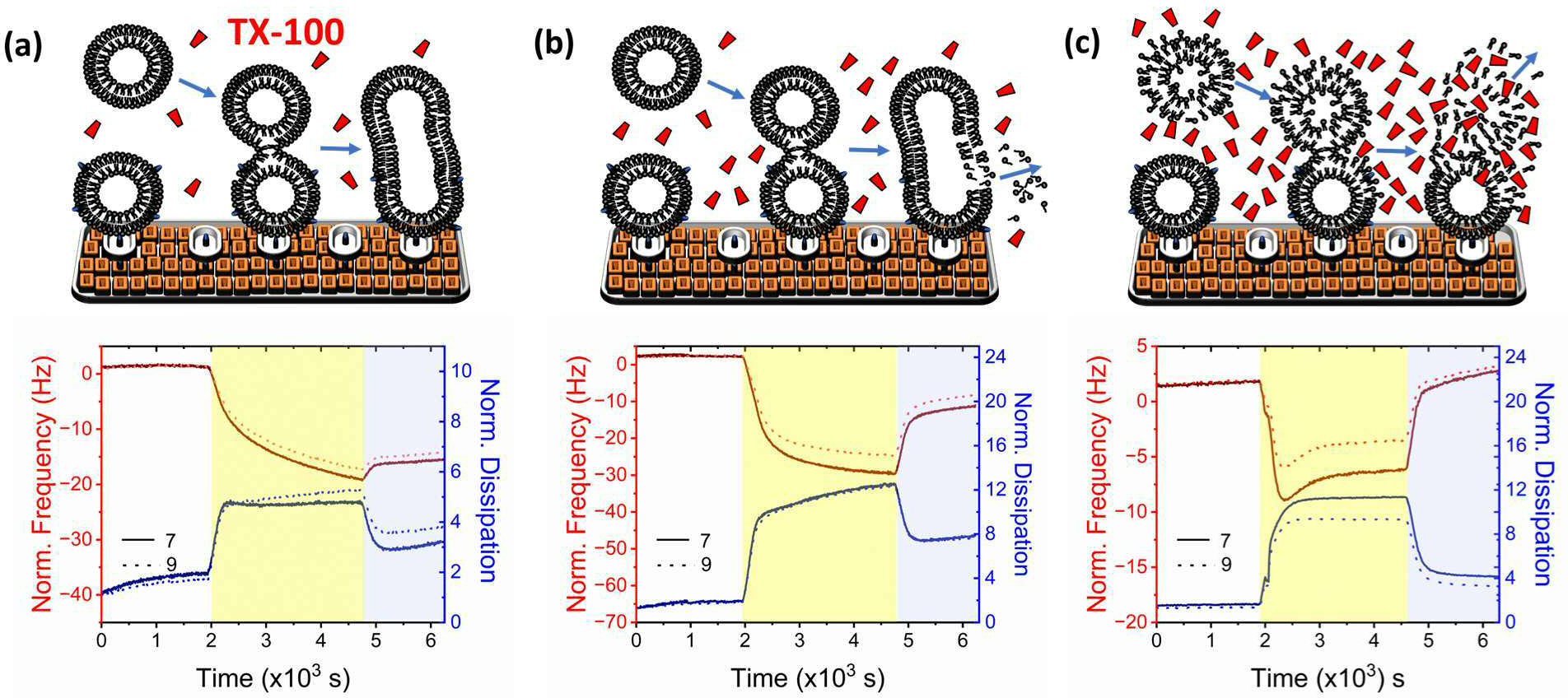
QCM-D analysis of TX-100 induced vesicle fusion on solid surfaces. Schematic illustrations (top panels) and normalized variation in frequency (red) and dissipation (blue) (bottom panels) associated with the 7^th^ (solid lines) and 9^th^ (dashed lines) harmonics upon injection of LUVs and (a) 0.1 mM, (b) 0.2 mM and (c) 0.3 mM TX-100 to a surface functionalized with BSA-biotin, NeutrAvidin and biotinylated LUVs. As illustrated, TX-100 induces a time-dependent increase in mass on the surface due to vesicles in solution merging with anchored vesicles, and minimal material is removed during surface washing when the TX-100 concentration is below the CMC. In contrast, tethered species lyse and dissociate when the detergent concentration exceeds the CMC. Injection of TX-100 and freely diffusing LUVs is denoted by the yellow shaded areas and was performed after washing the vesicle saturated surface with 50 mM Tris buffer (pH 8). The blue shaded areas correspond to the final wash step when post-interaction, the surface was rinsed with buffer. All experiments were performed at 21°C.

When the concentration of TX-100 was doubled, similar rate constants were observed (334 ± 4 s^-1^) for the frequency decay though we note that the magnitude of the relative frequency and dissipation responses increased 1.5- and 2.5-fold respectively pointing towards an abundance of fused states on the surface (**Figure 5b**). When both surfaces were washed with solution, only a partial recovery in ΔF and ΔD was observed indicating that most of the deposited material irreversibly attached to the immobilized layer (**Figures 5a and 5b**). However, when detergent concentrations above the CMC were in the injectant, the deposited mass returned towards the initial frequency and dissipation levels during the wash step, suggesting rupture of the freely diffusing vesicles upon interaction with the surface, and the subsequent removal of lipid mass (**Figure 5c**). Indeed, these assertions were further confirmed when variations in frequency were plotted against dissipation across the timescale of the fusion interaction (**Supplementary Figure 22**). The slope of the dissipation changes versus frequency shown in **Supplementary Figure 22** reflect variations in the viscoelastic properties of the immobilized layer across time. For detergent concentrations below the CMC, these changes broadly indicate that mass gain coupled with viscoelasticity enhancements on the surface precede minor levels of mass loss upon washing. At concentrations above the CMC, the vesicles in solution are susceptible to lysis and dissociation: a finding that we note is in line with our previous ensemble FRET approaches.

## Discussion

Several studies provide support for the hypothesis that TX-100 induces fusion of sub-micron sized vesicles. For instance, optical microscopy experiments have almost exclusively demonstrated GUV fusion to fluorescently-tagged native membranes at sub-solubilizing TX-100 concentrations^23^ and similar approaches have been employed to achieve TX-100-induced fusion between proteoliposomes and native membranes, resulting in the reconstitution of transmembrane proteins into GUVs^7, 8, 24^. In confocal-based fluorescence experiments, fusion was observed between permeable GUVs, suggestive that the process can occur through an initial contact between pores on adjoining vesicles^17^. While the mechanisms of GUV-versus LUV-detergent interactions have yet to be fully worked out, previous work indicates bilayer fluctuations, solubilization kinetics, detergent concentration requirements and detergent packing densities depend on the vesicle size and curvature^14, 25, 30, 53^. Thus, the development of techniques to dissect the multi-step mechanism of membrane fusion, especially at the higher end of the membrane curvature space are required.

To overcome this challenge, we implemented a robust in vitro fusion assay based on the measurement of energy transfer between lipophilic membrane dyes during lipid mixing. While FRET-based lipid mixing assays are not without shortcomings^54^ (for example, our observation that DiI is initially self-quenched suggests absolute FRET values may be underestimated and caution should be taken for accurate distance determinations) we demonstrate that the assay unveils multiple fusion intermediates as recorded by changes in the FRET efficiency and support our observations by a range of ensemble and single-particle measurements. Indeed, the cumulative data strongly supports vesicle fusion over alternative processes such as lipid exchange, evident by the marked increase in particle dimensions observed via DLS, SEM and Cryo-EM. Under symmetric TX-100 induced lipid exchange and dynamic equilibrium, the vesicle size distribution should remain relatively constant, but our experimental observations reveal significant particle size variations, pointing towards a fusion mechanism as the dominant process rather than lipid exchange. We do however acknowledge that we cannot completely exclude the possibility that some degree of POPC lipid exchange occurs, especially as vesicles in close proximity begin to interact. In this context, molecular dynamics simulations probing the interaction may be valuable. Nevertheless, the specific detergent employed and the speed through which the detergent leads to fusion and solubilization, categorized as “fast” or “slow” based on the detergent’s flip-flop rate are highly likely to govern the underlying mechanism^15^. In instances of slow solubilization, the process involves lipid extraction and/or the generation of micelles that are subsequently released and exchanged with enclosed vesicles. Conversely, in the context of TX-100, the rapid flip flop rate leads to the equal distribution of detergent molecules across both membrane leaflets, saturation of the membrane surface, and a pathway towards solubilization that occurs via open vesicular intermediates, which could facilitate fusion^25^. Here, examples of fused species induced by TX-100 were visualized by SEM and Cryo-TEM, both of which provide information on the morphology of the intermediates, and their occurrence under varying TX-100 conditions, and the combination of experiments strongly implies that the fusion process is unidirectional, precedes solubilization, and occurs via a mechanism that involves permeabilization, docking, hemifusion, and full lipid mixing. This is striking for two reasons: first, the fusion process occurs at sub-solubilizing detergent concentrations below the CMC and second, the fusion products observed here are remarkably similar to those observed during protein-induced vesicle fusion and viral fusion, raising the tantalising possibility that the observed intermediates are common, and share structural similarities, regardless of the fusogen.

A direct quantitative comparison of our findings on LUVs to previous studies on both LUVs and GUVs is challenging due to differences in membrane composition, vesicle size, the detergent-to-lipid ratio used and the precise measurement method. Nevertheless, it is clear the fused LUVs studied here, and those studied previously^13, 17, 23, 55^, share common attributes. This includes similar TX-100 concentration requirements to achieve fusion, observations of particle size and surface area enlargements upon TX-100 addition below the CMC, and vesicle permeabilization (which may suggest that TX-100 induced pores on adjacent vesicles facilitate fusion). Moreover our observation of nm-sized mixed detergent-lipid micelles under solubilizing conditions is in line with previous findings. Our studies on highly curved sub-micron sized vesicles are therefore complementary to previous optical microscopy and light scattering measurements and open a platform for exploring a matrix of environmental variables that may regulate the interaction.

Remarkably, our QCM-D results also show discrete steps of vesicle fusion and serve to highlight the benefits of using TX-100 to controllably fuse vesicles to surfaces containing immobilized vesicles: an important observation that may find direct relevance in a wide range of biotechnological applications. For example, in targeted drug delivery, vesicles encapsulating therapeutic agents are required to fuse to target sites, controllably releasing the encapsulated molecule. Vesicle fusion to surfaces can also be employed to create biomimetic systems that mimic natural cellular structures, or controllably modify and functionalize materials for enhancement of properties including conductivity and chemical reactivity. There is also scope for the fusion products to be used or manipulated in the context of sensor development, for instance in the detection of specific analytes, or in microfluidic devices where vesicle fusion can be employed to create well-defined membrane structures or facilitate the controllable transport and manipulation of small liquid volumes. We previously suggested that TX-100, along with other non-ionic surfactants induce swelling of lipid vesicles, but the absence of fusion observed with detergents such as Tween-20, suggest that the fusion efficacy is detergent-dependent (**Supplementary Figure 23**). Nevertheless, we expect the presented tools and techniques will be used to further define the fusion mechanism when different vesicle subpopulations are introduced, and to disentangle the impact of vesicle curvature, presence of lipid rafts and other environmental factors (pH, ionic strength) which may regulate the interaction. Indeed, the combined approaches and developed tools unambiguously dissect the underlying mechanism of TX-100-induced vesicle fusion and could become an indispensable toolbox for further characterizing the influence of fusogens, including those with important biomedical significance. Moreover, dissecting the mechanism of LUV fusion is critical, not only for expanding our fundamental knowledge of vesicle biology, but for the development of biocompatible and tissue-specific delivery systems.

A major aspect of our results that deserves attention is that sub-solubilizing concentrations of TX-100 were required to produce the fusion states described. While techniques such as isothermal titration calorimetry have been applied extensively to quantify the thermodynamics of detergent binding and partitioning below the CMC^12, 22, 56, 57^, our work, which quantifies single vesicle morphologies, implies that individual TX-100 monomers play a key role in initiating vesicle fusion. This is particularly remarkable because non-ionic detergents typically achieve structural changes only once above their CMCs. While further work is required to elucidate the precise role of the molecule in initiating fusion, we hypothesise that sufficiently high local detergent densities on the membrane may act as a nucleation site for docking, potentially via a pathway that involves creation of pores. Our approaches enabled fusion intermediates to be observed up to detergent: lipid ratios of ∼ 5: 1, suggesting that morphological rearrangements and solution exchange are initiated during LUV-detergent saturation. While membrane saturation and insertion of TX-100 into the bilayer may lead to local invaginations and permeabilization within intact vesicles prior to fusion, it is possible that bilayer bending due to vesicle swelling could also facilitate the process. It is also worth re-emphasizing a key benefit of the FRET-based strategy: unlike conventional phase contrast or fluorescence-based imaging, where only a 2-dimensional plane is captured, the FRET response reports on the dye separation distance across the entire 3-dimensional volume of the fused species.

## Conclusions

The approaches described here provide a general framework for dissecting the precise molecular level events that lead to TX-100 induced LUV fusion in vitro. TX-100 dynamically alters the conformation and integrity of both freely diffusing and surface-immobilized vesicles via a mechanism involving vesicle permeabilization, docking, hemifusion, and full lipid mixing, prior to solubilization and lysis. These observations provide mechanistic insights for how the widely used TX-100 detergent dynamically perturbs and modulates highly curved membrane structures, even at concentrations below the CMC. We anticipate that our current demonstration will be a starting point for addressing many important aspects of vesicle fusion, including but not limited to, real-time analysis of single-vesicle fusion events, and the influence of environmental factors, including cholesterol and lipid content, and may be directly relevant to key biotechnological applications where targeted vesicle fusion, especially to surfaces, is critical. We also expect that the developed approaches will find general utility for unveiling the multi-step mechanism of vesicle fusion in response to additional perturbative agents, including membrane-disrupting proteins and anti-viral agents, and could have important implications for evaluating drug delivery efficiencies.

## Methods

### Materials

1-palmitoyl-2-oleoyl-glycero-3-phosphocholine (POPC) and 1-oleoyl-2-(12-biotinyl(aminododecanoyl))-sn-glycero-3-phosphoethanolamine (biotin-PE) lipids in chloroform were purchased from Avanti Polar Lipids and used without additional purification. TX-100 was purchased from Sigma Aldrich and freshly suspended in 50 mM Tris buffer (pH 8) prior to each use. 1,1‘-Dioctadecyl-3,3,3‘,3‘-Tetramethylindocarbocyanine (DiI) and 1,1‘-Dioctadecyl-3,3,3‘,3‘-Tetramethylindodicarbocyanine (DiD)) were purchased from ThermoFisher Scientific and used without additional purification. All lipid stock solutions were stored in chloroform at −20°C prior to use, whereas DiI and DiD stock solutions from the manufacturer were stored at 4°C. Lipo-Cy3-CO and Lipo-Cy5-N_3_ were prepared as described elsewhere^38^.

### Preparation of vesicles incorporating DiI and DiD

Large unilamellar vesicles composed of lipid, biotin-PE and either DiI or DiD were prepared immediately prior to each experiment by extrusion. Briefly, unlabelled lipids and fluorescently labelled lipids were mixed in chloroform at the ratios specified in the main text prior to evaporation of the solvent by application of a gentle nitrogen stream. The resulting lipid film was resuspended in 50 mM Tris buffer (pH 8) and mixed well by vortex. Final solutions were then extruded through a polycarbonate membrane filter with size cut off of 200 nm prior to use. The same procedure was used to incorporate Lipo-Cy3-CO and Lipo-Cy5-N_3_.

### Preparation and purification of vesicles encapsulating Cal-520 and Ca^2+^

Vesicles encapsulating either Cal-520 or Ca^2+^ were prepared as previously described^30^. Briefly, chloroform was removed from POPC lipid stocks under gentle nitrogen flow, and the resulting thin lipid film was dissolved and vortexed in 50 mM Tris buffer (pH 8) containing either 100 mM Cal-520 or 10 mM Ca^2+^. The solutions were then extruded at least 21 times through a 200 nm cut off polycarbonate membrane filter (Avanti Polar Lipids). Vesicles were then filtered using PD-10 desalting columns (Sigma Aldrich) to separate non-encapsulated molecules from the loaded vesicles.

### Fluorescence Correlation Spectroscopy

FCS experiments were performed using a Zeiss LSM 880 microscope equipped with a 514 nm excitation line and GaAsP detector. LUVs incorporating 99 % POPC and 0.1 % DiI were diluted in 50 mM Tris buffer (pH 8) to a final lipid concentration of 6 μM, pipetted onto a pre-cleaned microscope slide, and sandwiched to a 1.5 glass coverslip via silica. Typical excitation powers were 4 μW as measured at the sample plane. Correlation curves were fitted to an equation of the form

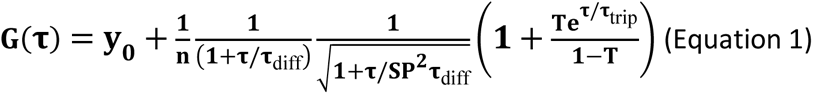

where *y***_0_**, *n*, *τ*, ***τ***_diff_, *SP*, *T* and ***τ****_trip_* are the offset, effective number of particles in the confocal volume, lag time, residence time in the confocal volume, structure parameter (which describes the shape of the detection volume and is equal to ***ω_xy_/ω_z_*** where ω is the width of the laser spot in the x, y and z directions), fraction of particles in the triplet state and residence time in triplet state, respectively. Diffusion coefficients were then determined and the hydrodynamic radius was estimated via the Stokes-Einstein relationship.

### Steady State FRET Vesicle Fusion Assay

200nm sized POPC vesicles incorporating 2 % DiI were suspended in 50 mM Tris buffer (pH 8) with those incorporating 2 % DiD at a DiI-LUV: DiD-LUV ratio of 1: 3 and final lipid concentration of 88 μM. The solution was then gently stirred by magnetic stirring at 1.5 Hz. Fluorescence emission spectra were then acquired using a HORIBA Fluoromax-4 spectrophotometer with excitation wavelengths of 520 nm and 620 nm. All emission spectra were corrected for background and acquired under magic angle conditions. Apparent FRET efficiencies were estimated via

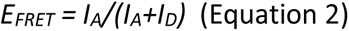

where *I_D_* (λ_ex_ = 520 nm; λ_em_ = 525 - 639 nm) and *I_A_* (λ_ex_ = 520 nm; λ_em_ = 640 - 800 nm) represent the integrated fluorescence emission intensities of DiI and DiD, respectively. We also measured the efficiency of energy transfer using the *RatioA* method which yields a value proportional to *E* by comparing the magnitude of sensitized acceptor emission after removal of donor bleedthrough to the emission obtained under direct excitation of the acceptor 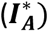 (λ_ex_ = 620 nm; λ_em_ = 640 - 800 nm). Here,

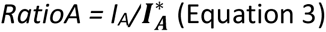

Plots of *E* versus *[TX-100]* are reported as the mean and standard error from three experimental runs.

### Time Correlated Single Photon Counting

Time-resolved fluorescence decays were acquired using a FluoTime 300 instrument equipped with a pulsed 530 nm excitation line (80 MHz) and PMA Hybrid 07 photomultiplier tube (Picoquant). Briefly, DiI and DiD loaded POPC vesicles were mixed and magnetically stirred at 1.5 Hz. Fluorescence lifetime decays at an emission wavelength of 565 nm were then collected under magic angle conditions until 10^4^ photon counts accumulated at the decay maximum. Decay curves were then fitted by iterative reconvolution of the instrument response function and the observed intensity decay, *I_t_*, using a multi-exponential decay of the form

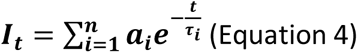

where *a_i_* and *t_i_* represent the fractional amplitudes and lifetimes of the i’th decay components. FRET efficiencies, *E*, were estimated via

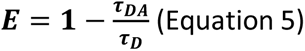

where τ*_DA_* and τ*_D_* represent the lifetime of DiI in the presence and absence of DiD, respectively. All experiments were performed in 50 mM Tris buffer (pH 8) using DiI-LUV: DiD-LUV ratios as specified in the main text and final lipid concentrations of 88 μM.

### Dynamic Light Scattering

Hydrodynamic radii of vesicles in TX-100-rich solutions were estimated using a Zetasizer mV DLS system (Malvern Panalytical) equipped with a 632.8 nm laser line. Briefly, POPC LUVs at a final lipid concentration of 88 μM were suspended in 50 mM Tris buffer (pH 8, n = 1.33) and TX-100 was added at the desired concentration. Correlation functions, g(τ), produced via the intensity of scattered light were fitted to a single-species model of the form

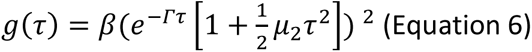

or a multi-species model of the form

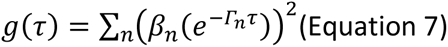

where *β* is the coherence factor, µ is the central moment of the distribution of decay rates and *Γ* is the decay rate. Diffusion times, τ, were then related to the diffusion coefficients, *D*, via

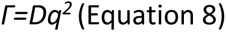

where *q* is the wave factor. Hydrodynamic radii were then extracted using the Stokes-Einstein equation as discussed elsewhere^58^. All DLS measurements were performed using 178° backscattering, and intensity-size distributions were fitted to a lognormal distribution of the form

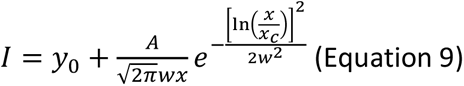

where *y_0_* is the offset, *x_c_* is the centre of the distribution, *w* is the log standard deviation and *A* is the area.

### Single Vesicle FRET Spectroscopy

Microfluidic flow cells were constructed^41^ and coated with 0.1 mg/mL BSA-Biotin, 1 mg/mL BSA and 0.2 mg/mL NeutrAvidin. Between each addition, the flow cells were washed with buffer to remove any unbound molecules. Biotinylated POPC vesicles incorporating 2 % DiD were mixed with non-biotinylated vesicles containing 2 % DiI (DiI-vesicle: DiD vesicle = 1: 3) at a final lipid concentration of 88 μM in TX-100 rich media (50 mM Tris buffer, pH 8). The fused species were then added to the functionalized coverslip for FRET-based imaging. Unbound vesicles were then removed by washing the flowcell with buffer. Objective-based TIRF microscopy was then performed on an inverted Nikon Eclipse Ti microscope containing a 100 x NA 1.49 oil immersion objective lens (Nikon)^41^. Excitation was achieved from a continuous wave 532 nm laser line (Obis, Coherent) with the diameter of the illuminated region in the field of view calculated to be 26 μm. To mitigate against photobleaching, typical excitation intensities were 8 μWcm^-2^. DiI and DiD emission was spatially separated using a DualView Image Splitter (Photometrics) containing a dichroic (T640LPXR, Chroma) and band pass filters (ET585/65M and ET700/75M, Chroma) and imaged in parallel on a back-illuminated Prime 95b CMOS camera cooled to −30°C (Photometrics). Fluorescence and FRET trajectories from immobilized vesicles were acquired with 50 ms exposure time and recorded images were analysed in MATLAB (R2019a) using iSMS FRET microscopy software^59^. Briefly, co-localized DiI and DiD emission trajectories, termed *I_D_* and *I_A_* respectively, were obtained by integration of the intensity within the area containing the vesicle of interest at each time point, and apparent FRET efficiencies, *E_FRET_*, were estimated as previously described. In line with previous work^28^, tri-Gaussian fits were applied to the distributions shown in **Figure 3d** and based on the crossing points of the individual components, thresholds for deciding if the species were docked (*E_FRET_* < 0.25), partially fused (0.25 ≥ *E_FRET_* ≤ 0.55) or fully fused (*E_FRET_* > 0.55) were used.

### Quartz Crystal Microbalance with Dissipation Monitoring

QCM-D experiments were performed using a Q-Sense E4 system (Biolin Scientific) instrument. Briefly, SiO_2_ coated substrates (Biolin Scientific) with a fundamental frequency of 5 MHz were first cleaned by UV ozone for 10 minutes, then sonicated in 2 % Hellmanex III for 10 minutes and ultrapure Milli-Q water for 20 minutes. The sensors were then dried under nitrogen before being treated with UV ozone for an additional 30 minutes. The sensors were then immersed in 4 % v/v ATES/IPA solution for 16 hours to produce an amine monolayer and dried under nitrogen before being installed in the sensor housing module. All sensors were washed with 50 mM Tris (pH 8) under a flow rate of 20 μL per minute until a stable baseline, defined as < 5 Hz shift per minute was achieved. At the start of each experiment, the sensor surfaces were functionalized sequentially with 0.1 mg mL^-1^ biotinylated bovine serum albumin (BSA), 1 mg mL^-1^ BSA and 0.2 mg mL^-1^ NeutrAvidin as per the single-vesicle TIRF experiments. The sensors were also washed with buffer after each addition to remove any unbound molecules. Vesicles incorporating 1 % Biotin-PE were then flushed across the surface at a final lipid concentration of 0.25 mg mL^-1^ until saturation of the surface was reached, followed by another buffer wash step. Next, POPC vesicles lacking Biotin-PE at a final lipid concentration of 0.16 mg mL^-1^ were mixed with TX-100 at the specified concentrations and immediately washed over the sensor surface. Unlike the TIRF measurements, here we monitored the fusion of vesicles in solution to vesicles pre-tethered to the sensor substrate.

### Scanning Electron Microscopy

SEM images were acquired using a JEOL JSM 7800-F system operating at 5kV. Vesicles were induced to fuse in 50 mM Tris (pH 8) conditions containing TX-100 as specified in the main text, diluted, and deposited onto a silicon substrate. The solution was then evaporated, and the immobilized vesicle layer was sputtered with a 5 nm layer of Pt/Pd. We note that the final concentration of vesicles applied to the substrate was sub-nM to prevent the formation of conglomerates that are not directly correlated to the effects of TX-100. This was confirmed by the fact that for low TX-100 concentrations, and for vesicles imaged in the absence of detergent, we did not observe any vesicle clusters. Vesicle diameters were determined using ImageJ, where automated analysis of black and white binary images enabled separation of regions of white pixels (vesicles) against a dark background. Vesicle circularity, ϕ, was measured via

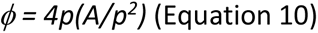

where *A* is the 2-dimensional surface area of the vesicle and *p* is the vesicle perimeter.

### Cryo-TEM

Cryo-TEM was used to directly visualize fused vesicle species^47^. Briefly, Quantifoil copper R 1.2/1.3 200 mesh grids (Electron Microscopy Sciences) were prepared by glow discharging at 20 mA and 0.26 mbar for 1 minute in a Pelco easiGlow glow discharge system. Small volumes of vesicles pre-incubated with TX-100 were than applied to the carbon side of the EM grid in 90 % humidity. Excess liquid was blotted off and the grids were plunge frozen into precooled liquid ethane using a Vitrobot system (Thermo Scientific), allowing fused species to be embedded and preserved within a thin layer of amorphous ice. Micrographs were then obtained using a Thermo Scientific Glacios Cryo-TEM microscope using an operating voltage of 200 kV and 120,000x magnification. Vesicle diameters and circularities were determined using in-house computational analyses.

## Acknowledgements

This work was supported by Alzheimer’s Research UK (RF2019-A-001) and EPSRC (EP/P030017/1, EP/W024063/1). We thank Prof. Daniella Barillá (Department of Biology, University of York) for use of DLS instrumentation, Dr. Jamie Blaza (Department of Biology, University of York) for technical expertise and use of Cryo-TEM, and Prof. Thomas Krauss (University of York) for use of SEM facilities. We also thank the Bioscience Technology Facility (University of York) for use of FCS and fluorescence spectroscopy facilities, and Dr Laurence Wilson (University of York) for critically reading the manuscript. The authors also thank Dr. Christopher D. Spicer and Lydia J. Barber for helpful discussions and for the generous donation of Lipo-Cy3-CO, Lipo-Cy5-N_3_ and dialysis kits. We also thank the peer reviewers for critically evaluating our manuscript and for providing positive and constructive comments.

## Data Availability

The datasets generated during the current study are available from the corresponding author on reasonable request.

## Author Contributions

S. D. Q. designed the research and wrote the manuscript with the help of all other co-authors. L. G. D., C. K., D. C., C. M. H., N. S, and S. D. Q. collected the data and performed the research. L. G. D., C. K., D. C., C. M. H., N. S., L. C., J. C. P., M. C. L., and S. D. Q. analyzed the data. S. D. Q. was the project administrator.

