## Supplementary Information for "Multiple Intermediates in the Detergent-Induced Fusion of Lipid Vesicles"

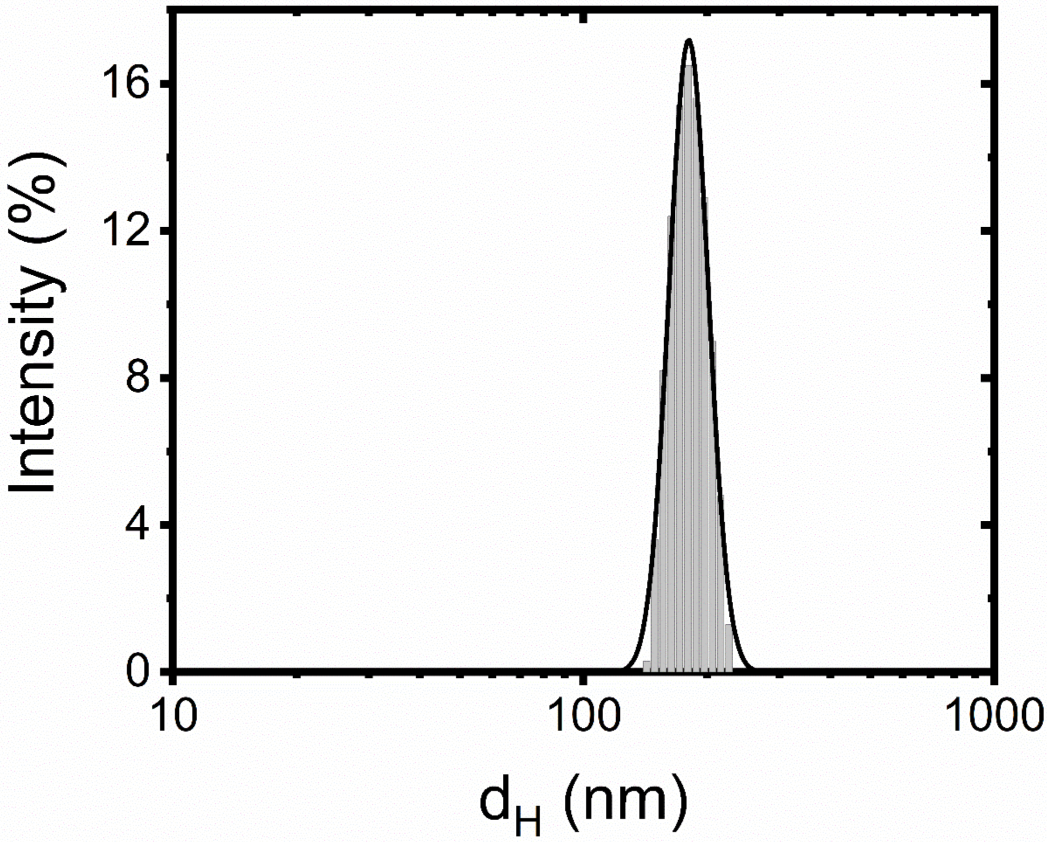


**Supplementary Figure 1. Representative hydrodynamic diameter ($d_H$) distribution of extruded POPC LUVs at 21°C obtained via dynamic light scattering**. The solid black line represents a lognormal fit with peak $d_H$ of 182.6 ± 0.1 nm ($\chi_r^2$ = 0.02).

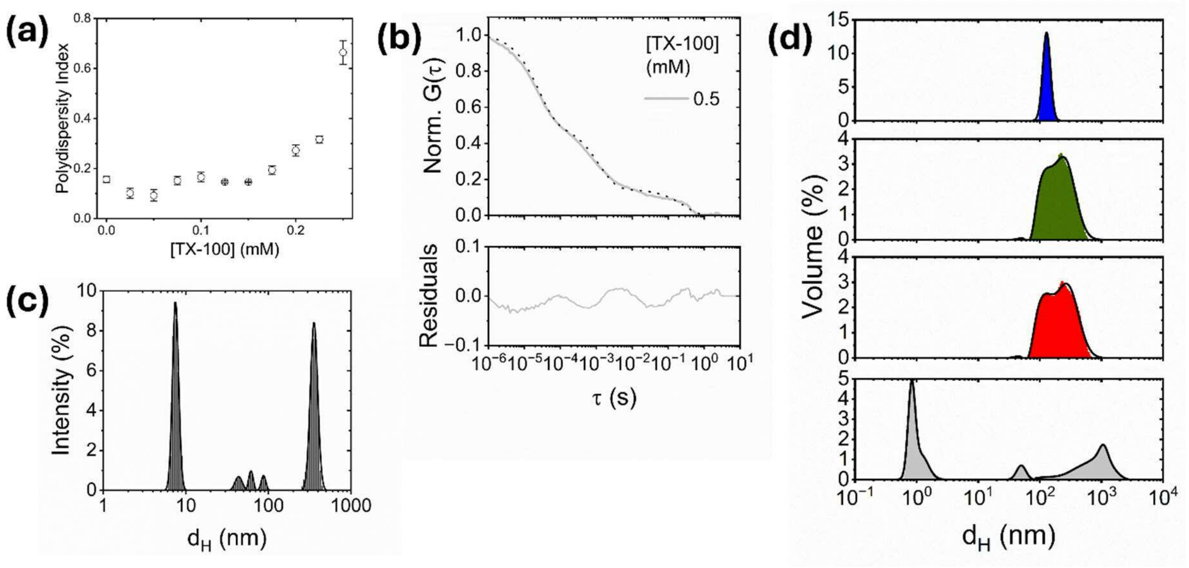


**Supplementary Figure 2. Characterization of vesicle-detergent interactions by DLS.** (a) Variation in DLS polydispersity index obtained from POPC LUVs in TX-100 rich solution. Data points and error bars represent the mean and standard error of the mean from three experimental runs. (b) Normalized variation in DLS correlation curve (solid line) and fit (dotted line) associated with interactions between LUVs and 0.5 mM TX-100. Bottom: residuals of the fit. (c) Hydrodynamic diameter versus intensity distribution obtained from LUVs in the presence of 0.5 mM TX-100. (d) Representative volume distributions obtained from POPC LUVs in the absence (blue) and presence of 0.15 mM (green), 0.2 mM (red) and 0.5 mM TX-100 (grey). All experiments were conducted at 21°C.

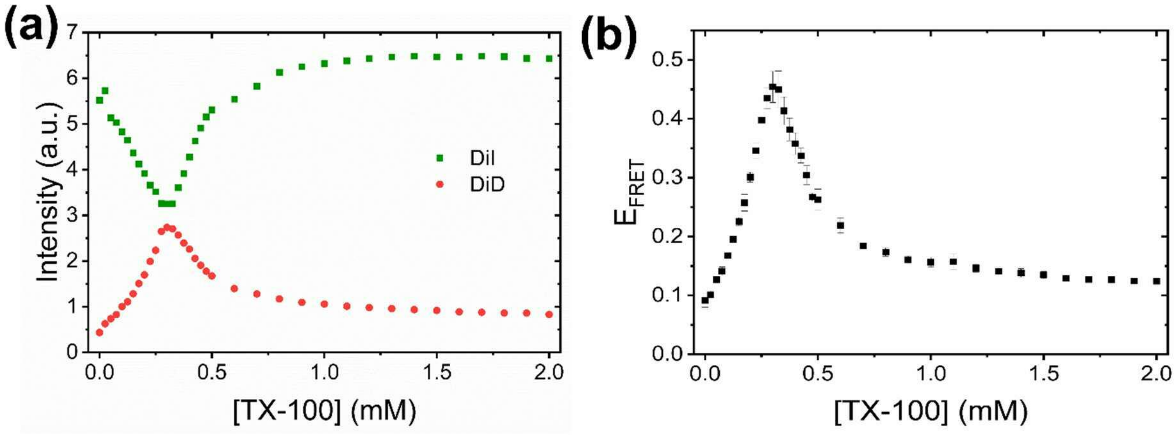


**Supplementary Figure 3. TX-100 induces lipid mixing between freely diffusing vesicles.** Representative variation in DiI and DiD fluorescence emission intensities and (b) apparent FRET efficiency ($E_{FRET}$) across the titration. Error bars represent the standard error of the mean obtained from three experimental runs. Solution conditions: 50 mM Tris, pH 8.0, [POPC] = 88 μM, 21$^{o}$C.

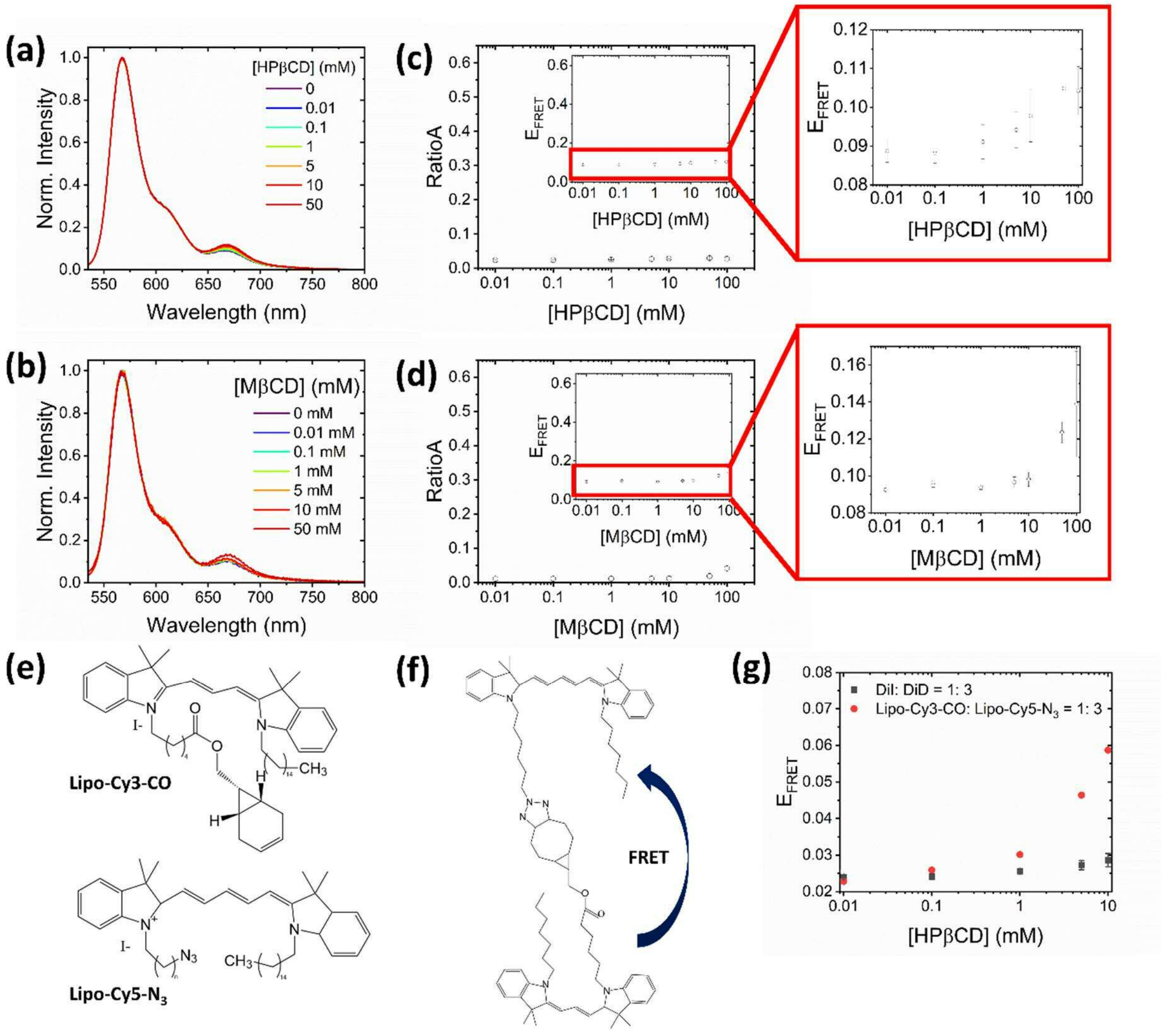

**Supplementary Figure 4. FRET response obtained from POPC LUVs in the presence of HPβCD and MβCD.** Representative variation in the fluorescence emission spectra ($\lambda_{ex}$ = 520 nm) obtained from DiI and DiD LUVs in the absence and presence of (a) HPβCD and (b) MβCD. (c-d) The corresponding variations in *RatioA* and $E_{FRET}$ are also shown. (e) Chemical structures of Lipo-Cy3-CO and Cipo-Cy5-$N_3$ and (f) upon cycloaddition. (g) The corresponding variation in $E_{FRET}$ recorded from LUVs containing Lipo-Cy3-CO and Lipo-Cy5-$N_3$ upon addition of HPβCD. Error bars represent the standard error of the mean from three separated experimental runs. Solution conditions: [POPC] = 88μM, Donor: Acceptor = 1: 3, 50 mM Tris, pH 8.0, 21$^{o}$C.

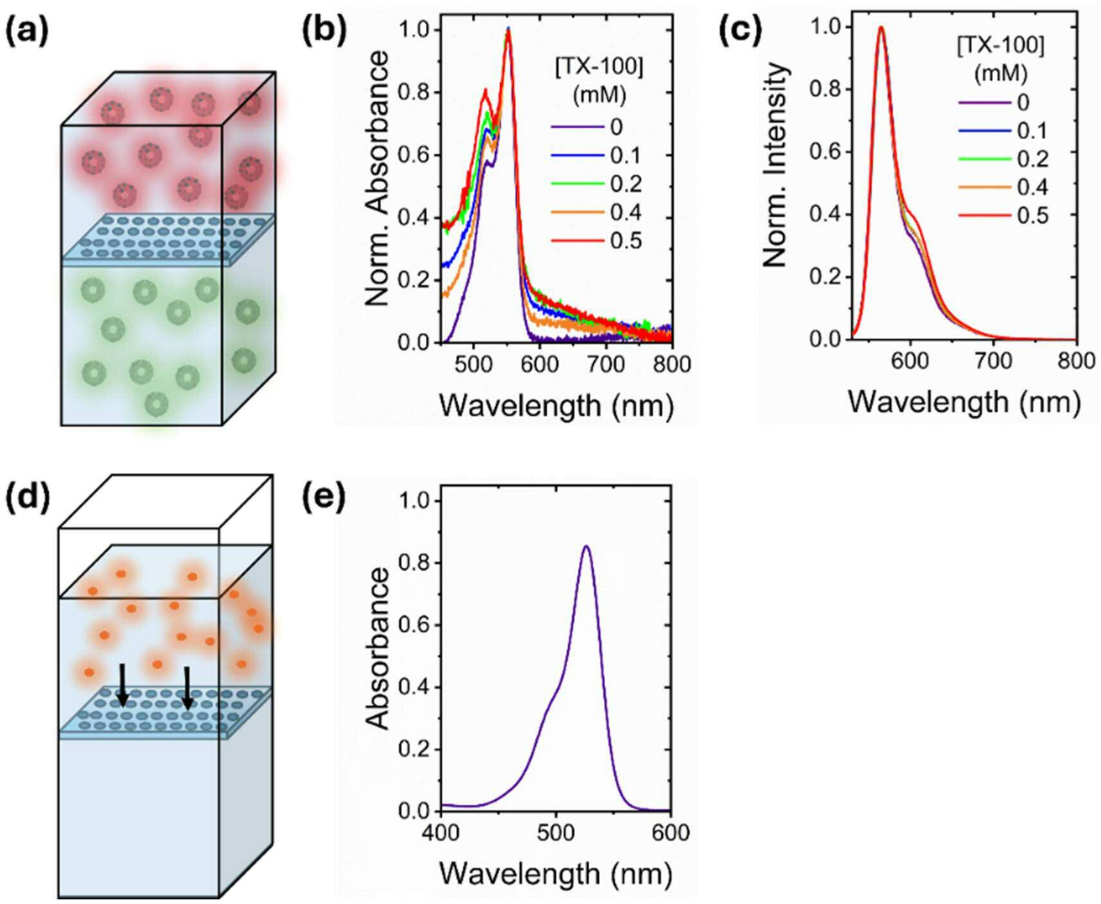


**Supplementary Figure 5. TX-100 does not induce lipid exchange.** (a) Schematic illustration of the lipid exchange assay. POPC LUVs containing 2 % DiI (green) and 2 % DiD (red) (1: 3) are separated by a 3 kDa MWCO dialysis membrane. After addition of TX-100 and incubation, lipid exchange was assessed by evaluating the absorption and fluorescence emission spectra of the solutions. (b) Representative absorption spectra obtained from the DiI solution in the absence and presence of TX-100, and after 5 hours. (c) The corresponding variation in fluorescence emission spectra ($\lambda_{ex}$ = 520 nm). (d) Illustration of the single dye exchange assay. A solution containing Rh6G was separated from clean buffer solution by a 3 kDa MWCO dialysis membrane. After 5 hours incubation, dye exchange was assessed by evaluating (e) the absorption spectra of the solution on the opposite side of the membrane, confirming dye exchange over the timescale of the experiment.

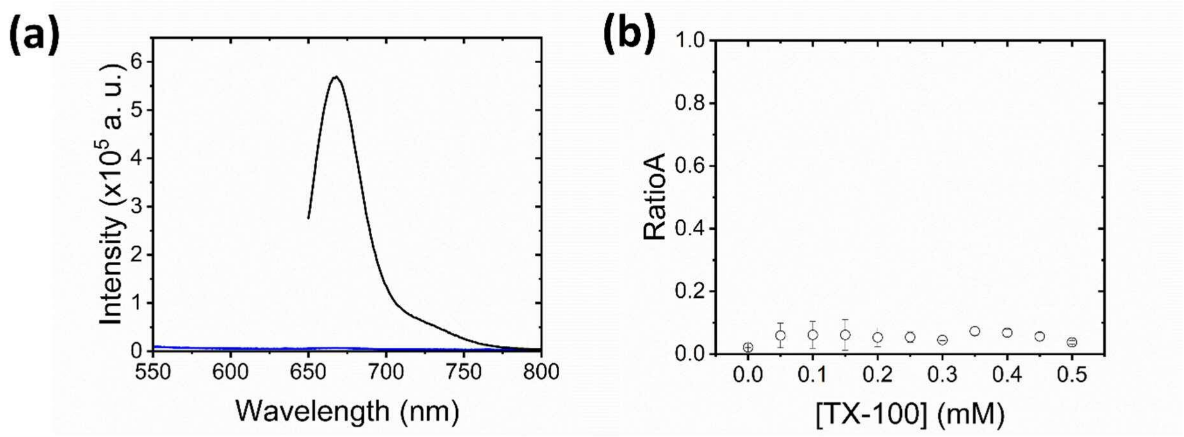


**Supplementary Figure 6. *RatioA* response obtained from DiD-labelled POPC LUVs in the presence of TX-100.** (a) Representative variation in the fluorescence emission spectra ($\lambda_{ex}$ = 520 nm (blue) and $\lambda_{ex}$ = 640 nm (black)) obtained from unlabelled LUVs incubated in the presence of those containing 2 % DiD in the presence of 0.25 mM TX-100. (b) Variations in the observed *RatioA* across the TX-100 titration. Error bars represent the standard error of the mean from three separated experimental runs. Solution conditions: [POPC] = 88μM, unlabelled: DiD LUVs = 1: 3, 50 mM Tris, pH 8.0, 21$^{o}$C.

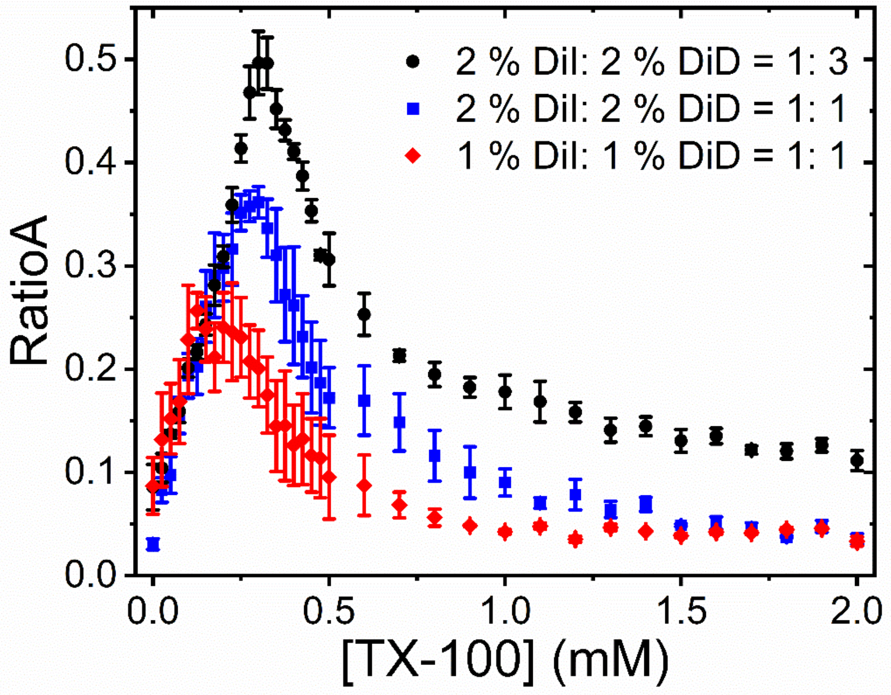


**Supplementary Figure 7. Optimization of the FRET-based lipid mixing assay.** Representative variation in the efficiency of resonance energy transfer, measured via the *RatioA* parameter, shown for solutions containing (i) 1 % DiI- and 1 % DiD-loaded vesicles incubated at a 1: 1 ratio (red) (ii) 2 % DiI- and 2 % DiD-loaded vesicles incubated at 1: 1 ratio (blue), and (iii) 2 % DiI- and 2 % DiD-loaded vesicles incubated at a 1: 3 ratio (black). Data points and error bars represent the mean and standard error of the mean obtained from three experimental runs. Solution conditions: 50 mM Tris, pH 8.0, [POPC] = 88 μM, 21°C.

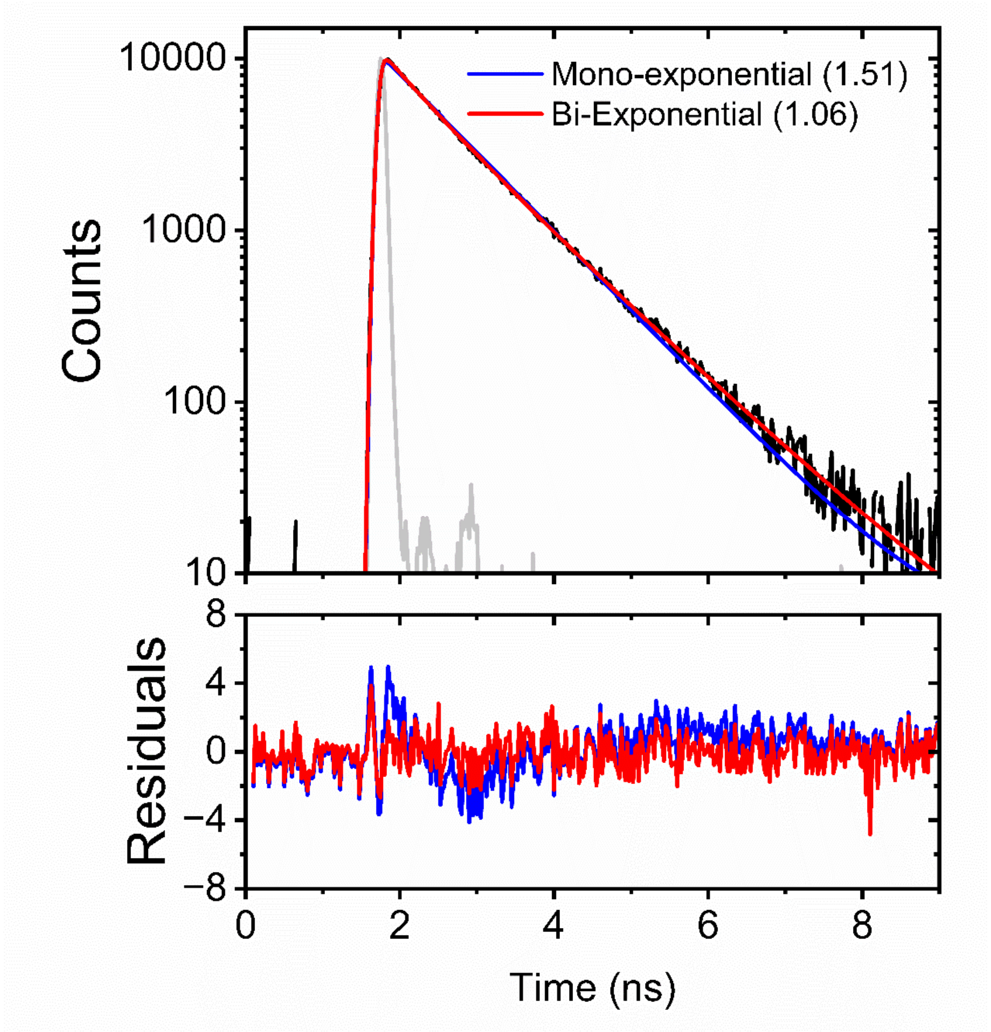


**Supplementary Figure 8. Time-resolved fluorescence decays obtained from DiI-loaded POPC fit to a bi-exponential decay model.** Top panel: Representative time-resolved fluorescence decay (black) obtained from POPC vesicles containing DiI in the presence of vesicles labelled with DiD in the absence of TX-100 and at a donor-vesicle: acceptor-vesicle ratio of 1:3. Solid blue and red lines represent fits to mono- and bi- exponential decay functions, respectively. The solid grey line represents the instrument response function (IRF). The numbers in brackets represent $\chi 2$ values. Bottom panel: residuals obtained from application of mono- and bi-exponential fits to the experimental data. Solution conditions: 50 mM Tris, pH 8.0, [POPC] = 88 $\mu$M, 21$^{o}$C.

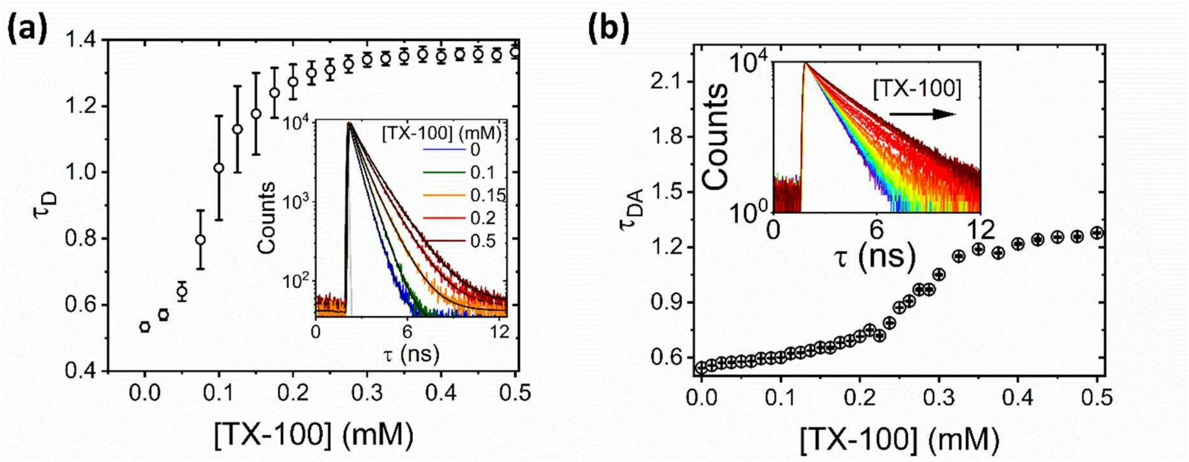


**Supplementary Figure 9. Time-resolved fluorescence decays associated with DiI in the absence and presence of DiD.** Variation in $\tau_D$ and $\tau_{DA}$ obtained from 2 % DiI-loaded vesicles in the (a) absence and (b) presence of vesicles containing 2 % DiD. Insets show the corresponding fluorescence decays. Solution conditions: 50 mM Tris, pH 8.0, [POPC] = 88 μM, 21°C. Data points represent the mean and standard error of the mean obtained from three experimental runs.

**Supplementary Table 1.** Fitting parameters and errors associated with bi-exponential fits applied to sensitized DiD fluorescence emission traces shown in **Figure 2f.**

| | 0.1 mM TX-100 | 0.2 mM TX-100 | 0.25 mM TX-100 |
|---|---|---|---|
| $Y_o$ | (11 ± 0.5) x10$^5$ | (25 ± 0.7) x10$^5$ | |
| $A_1$ | (40 ± 11) x10$^5$ | (44 ± 0.8) x10$^5$ | (28 ± 0.4) x10$^5$ |
| $t_1$ (s) | 30 ± 2 | 71 ± 1 | 225 ± 1 |
| $A_2$ | (53 ± 0.4) x10$^5$ | (16 ± 0.3) x10$^5$ | (48 ± 5) x10$^4$ |
| $t_2$ (s) | 1247 ± 24 | 1055 ± 11 | 716 ± 30 |
| $t_{average}$ (s) | 273 ± 6 | 897 ± 9 | 396 ± 11 |
| $R^2$ | 0.95 | 0.99 | 0.99 |

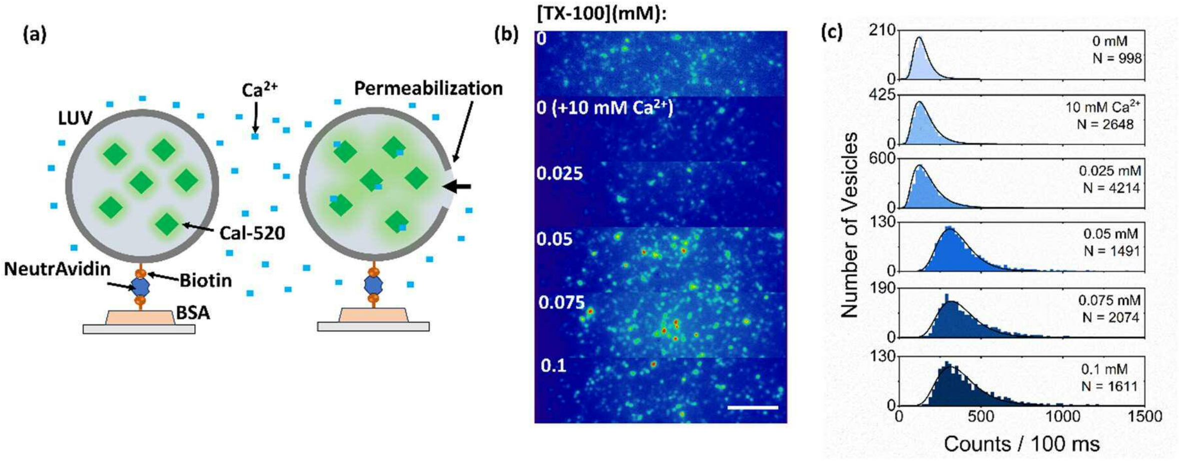


**Supplementary Figure 10. TX-100 induces vesicle permeabilization.** (a) Schematic illustration of the assay. LUVs containing Cal-520 are incubated in $Ca^{2+}$ buffer (left panel). Membrane permeabilization after TX-100 interaction allows $Ca^{2+}$ to enter the vesicle leading to a Cal-520 intensity enhancement (right panel). (b) Representative wide-field TIRF images of Cal-520 loaded vesicles in the absence and presence of TX-100. Scale bar = 5 μm. Note: all detergent-rich buffer solutions contained 10 mM $Ca^{2+}$. (c) Corresponding population intensity histograms obtained from N LUVs. The solid black lines represent log-normal fits to the experimental data.

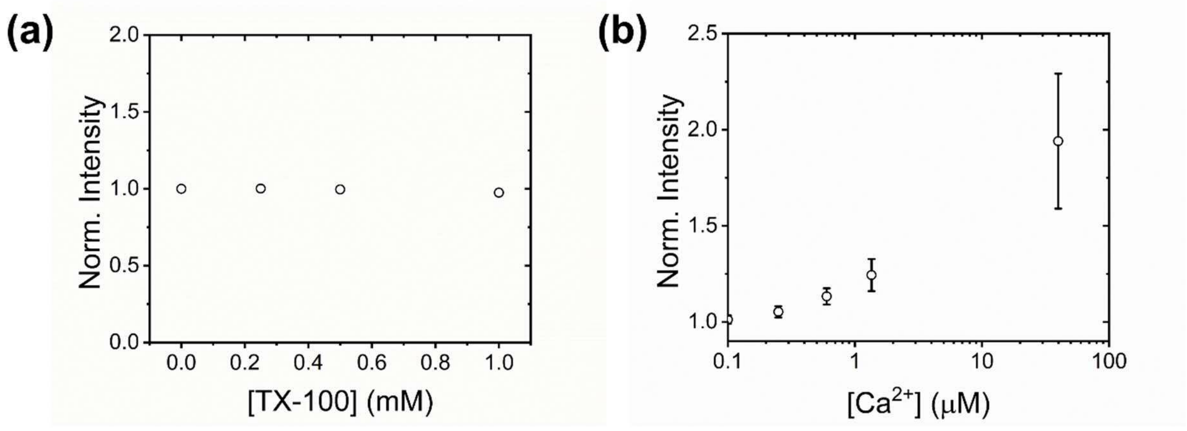


**Supplementary Figure 11. Representative variation in free Cal-520 emission intensity.** Normalized variation in the fluorescence emission intensity of 0.1 μM Cal-520 as (a) a function of TX-100 above and below the CMC, and (b) as a function of $Ca^{2+}$ in the presence of 0.5 mM TX-100 at 21°C. In both cases, the buffer solution was 50 mM Tris, pH 8.0, 21°C. Data points represent the mean and standard error of the mean obtained from three experimental runs.

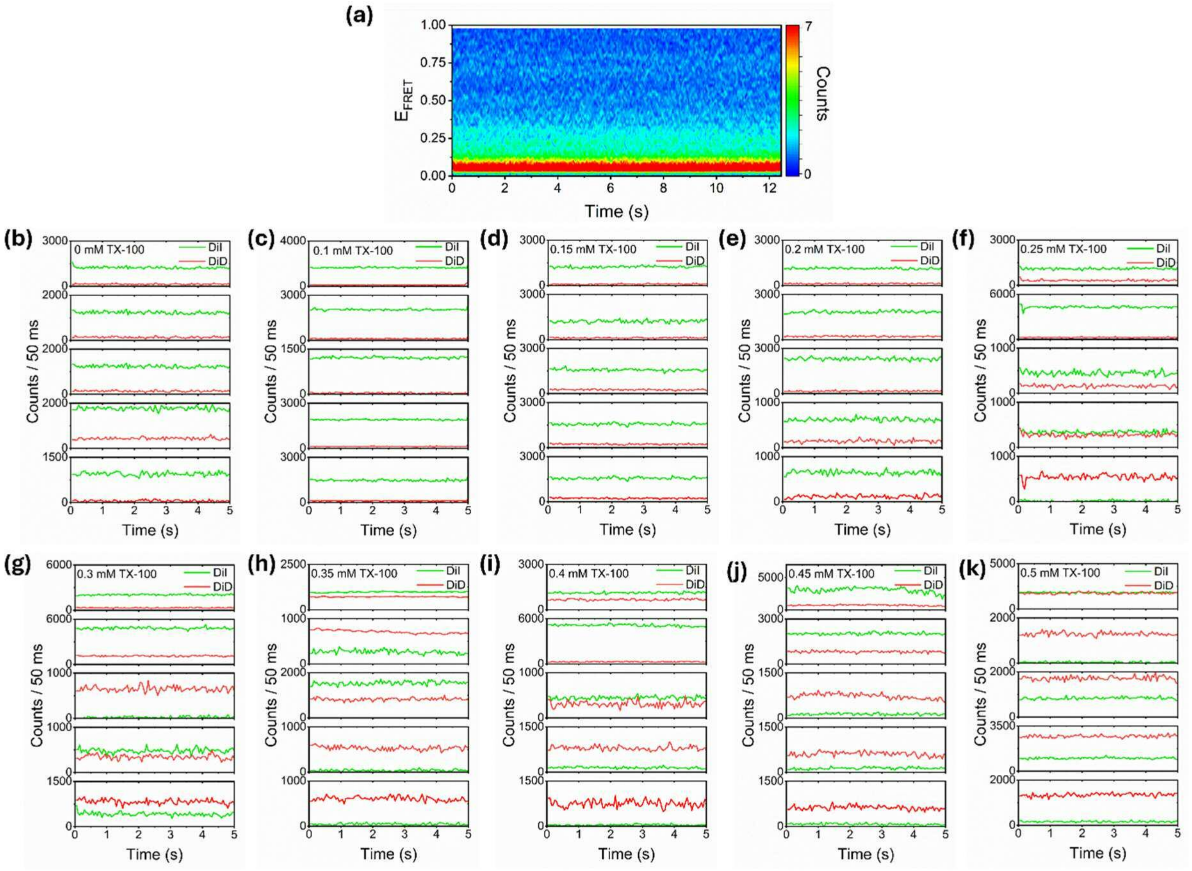


**Supplementary Figure 12. Stability of single-vesicle FRET trajectories.** (a) Representative contour plot of the time evolution of the FRET-active population (N = 50) in the absence of TX-100. Also shown are representative DiI (green) and DiD (red) intensity traces obtained in the (b) absence and (c-k) presence of TX-100. All data was collected at 21$^{o}$C.

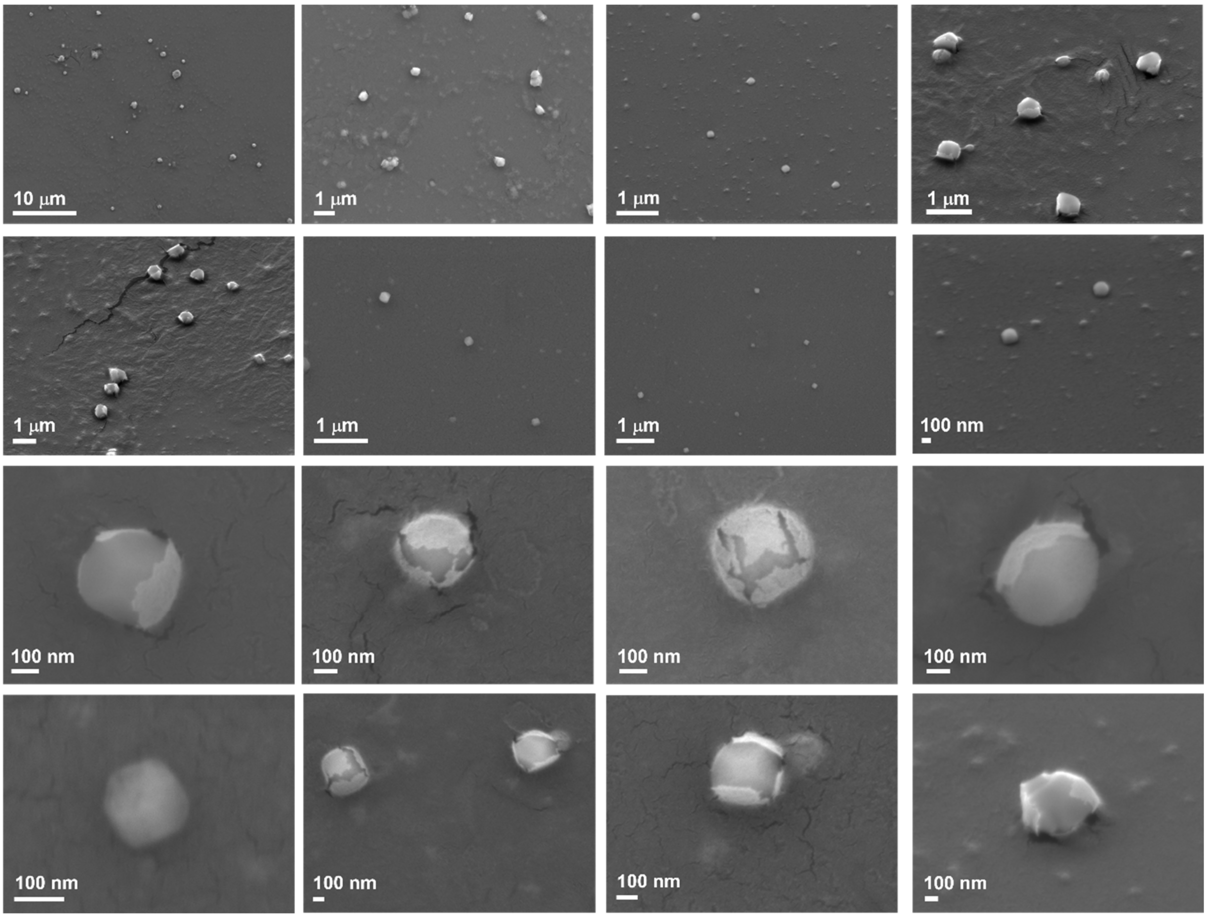


**Supplementary Figure 13. Representative SEM images of freshly prepared vesicles in the absence of TX-100.** Vesicles were prepared at 21ºC prior to SEM analysis.

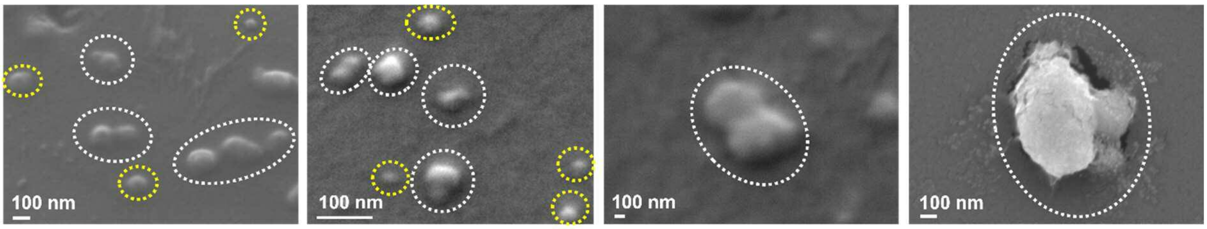


**Supplementary Figure 14. Representative SEM images of freshly prepared vesicles incubated with 0.15 mM TX-100.** Objects shown within the yellow and white dotted lines were classified as intact and docked, respectively. Vesicles were incubated with TX-100 at 21$^{o}$C prior to SEM analysis.

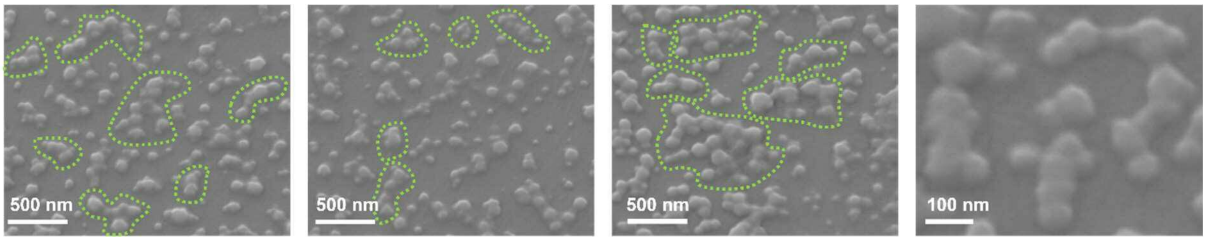


**Supplementary Figure 15. Representative SEM images of freshly prepared vesicles incubated with 0.15 mM TX-100.** Objects shown within the green dotted lines were classified as hemi-fused. Vesicles were incubated with TX-100 at 21$^{o}$C prior to SEM analysis.

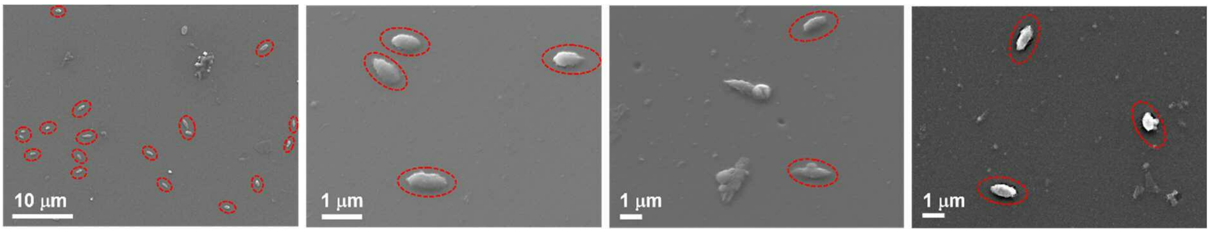


**Supplementary Figure 16. Representative SEM images of freshly prepared vesicles incubated with 0.15 mM TX-100.** Objects shown within the red dotted lines were classified as fully fused. Vesicles were incubated with TX-100 at 21$^{o}$C prior to SEM analysis.

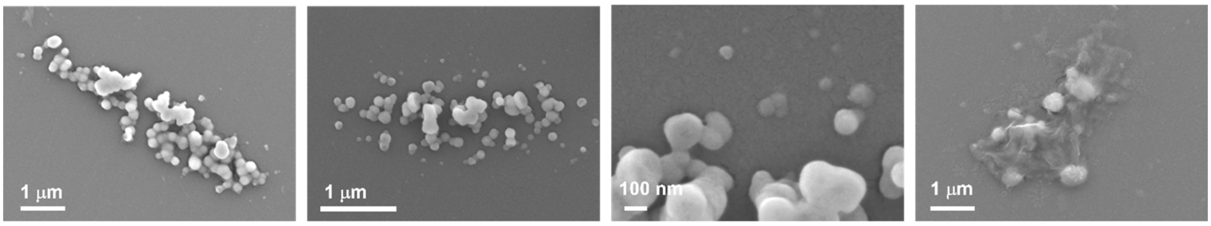


**Supplementary Figure 17. Representative SEM images of conglomerates obtained from incubating freshly prepared vesicles with 0.15 mM TX-100.** Vesicles were incubated with TX-100 at 21$^{o}$C prior to SEM analysis.

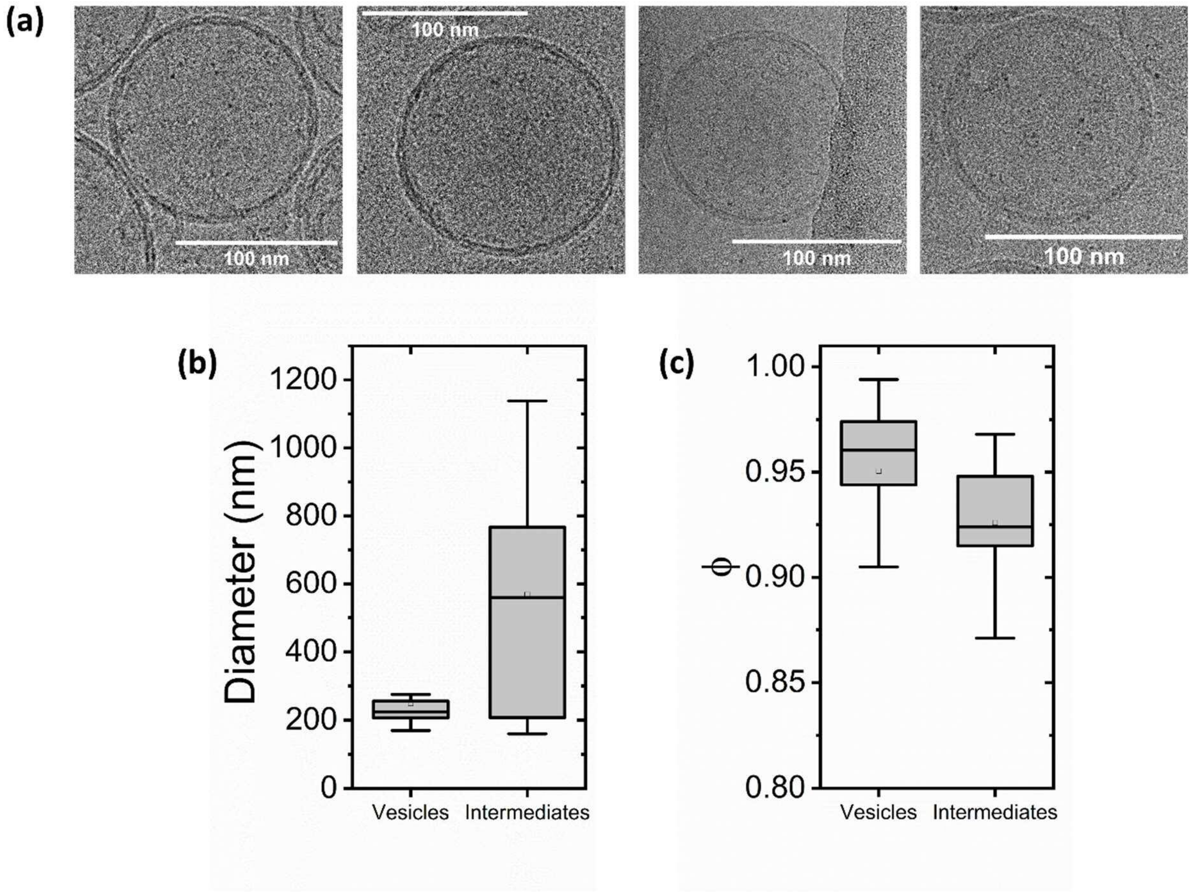


**Supplementary Figure 18. Characterization of POPC vesicles in the absence and presence of 0.15 mM TX-100 by cryo-EM.** (a) Representative cryo-TEM images of unilamellar vesicles obtained in the absence of TX-100. Comparative (b) size and (c) circularity distributions obtained from intact vesicles and fusion intermediates in the presence of 0.15 mM TX-100. The boxes represent the interquartile ranges, the upper and lower whiskers represent the 5$^{th}$ and 95$^{th}$ percentiles, the solid lines inside the boxes represent the medians, and the square represents the mean. Vesicles were prepared at 21$^{o}$C prior to Cryo-TEM analysis.

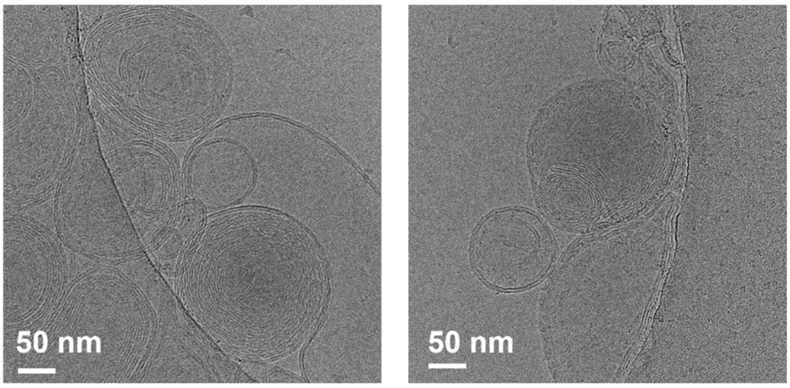


**Supplementary Figure 19. Representative Cryo-TEM images of multilamellar vesicles obtained in the absence of TX-100.** Vesicles were prepared at 21$^{o}$C prior to Cryo-TEM analysis.

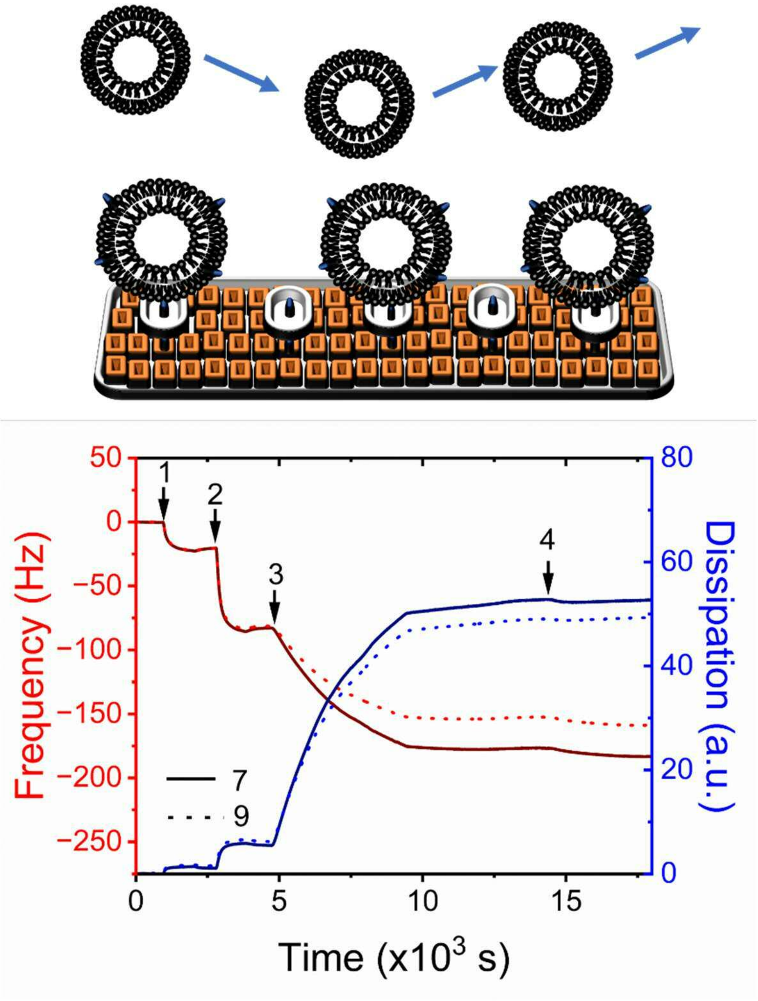


**Supplementary Figure 20. QCM-D of biotinylated vesicle immobilization and interactions with freely diffusing vesicles.** Evolution of the frequency (red) and dissipation (blue) responses (7$^{th}$ and 9$^{th}$ harmonics) upon the addition of (1) BSA-Biotin, (2) NeutrAvidin and (3) biotinylated LUVs to a $SiO_2$ substrate at 21$^{o}$C. After vesicle saturation, the sensor was rinsed with buffer ($10^3$s) before a solution of (4) non-biotinylated vesicles (final lipid concentration = 0.16 mg/mL) was flushed over the surface as schematically shown.

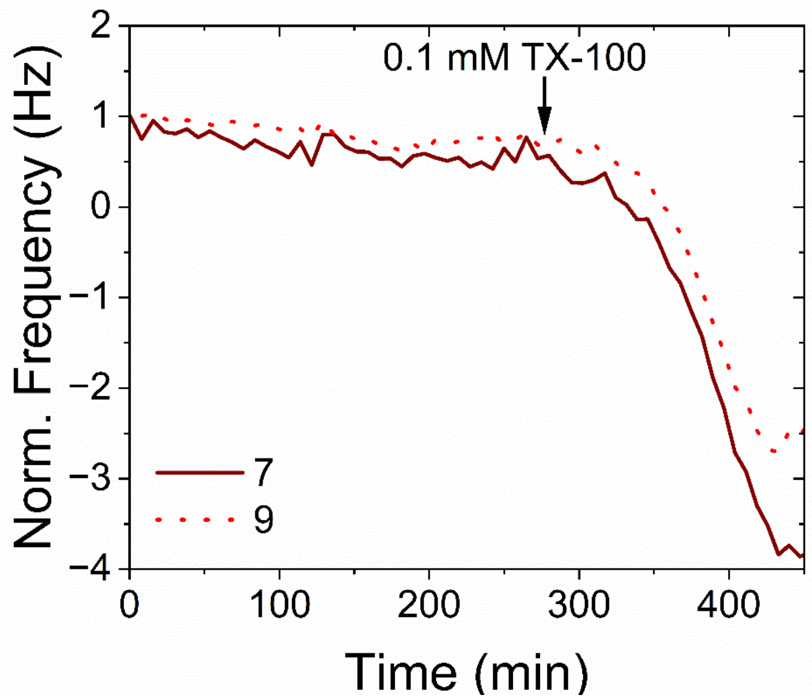


**Supplementary Figure 21. QCM-D of interactions between surface-immobilized vesicles and 0.1 mM TX-100 in solution.** Normalized variation in the frequency response of the 7$^{th}$ and 9$^{th}$ harmonics upon the addition of 0.1 mM TX-100 to biotinylated LUVs immobilized to a $SiO_2$ substrate at 21°C via biotin-Avidin interactions. The black arrow indicates the point of TX-100 injection.

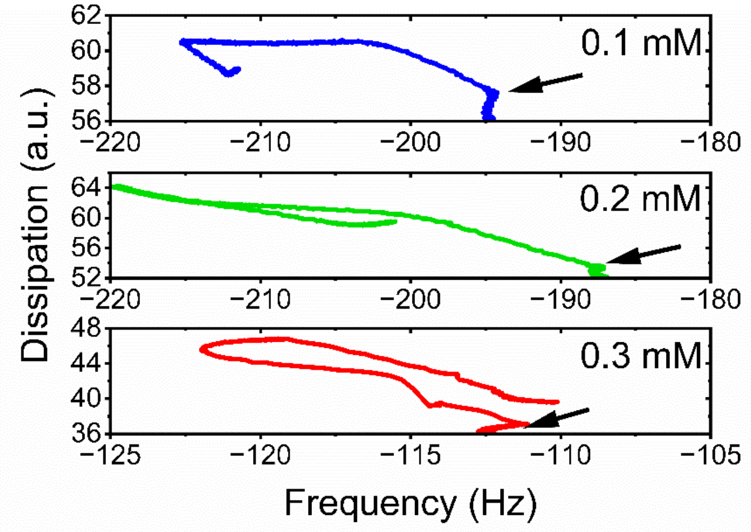


**Supplementary Figure 22. QCM-D of interactions between surface-immobilized vesicles, freely diffusing vesicles, and TX-100.** Frequency versus dissipation observed during the interaction between surface immobilized vesicles and 0.1 mM (top panel), 0.2 mM (middle panel) and 0.3 mM (bottom panel) TX-100 and POPC vesicles at a final lipid concentration of 0.16 mg/mL. The arrows indicate the point at which TX-100 and LUVs were injected. All experiments were performed at 21°C.

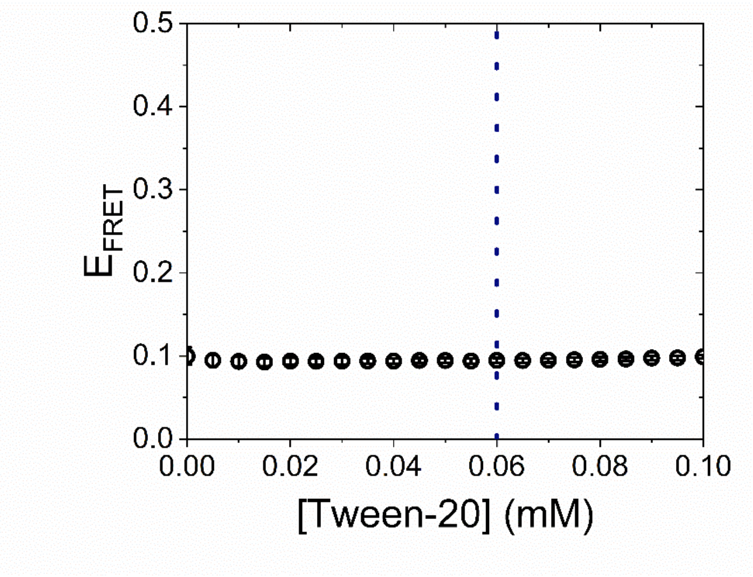


**Supplementary Figure 23. FRET-based lipid mixing assay in the presence of the non-ionic detergent Tween-20.** Representative variation in the apparent efficiency of resonance energy transfer, shown for a solution containing POPC vesicles in the absence and presence of Tween-20. Data points represent the mean and standard error of the mean obtained from three experimental runs. Vesicles contained 2 % DiI- and 2 % DiD and were incubated at a 1: 3 ratio. Solution conditions: 50 mM Tris, pH 8.0, [POPC] = 88 μM, 21°C. The dashed line corresponds to the CMC of Tween 20 (~0.06 mM).
